# Acoustically evoked K-complexes together with sleep spindles boost verbal declarative memory consolidation in healthy adults

**DOI:** 10.1101/2023.06.29.546822

**Authors:** Sven Leach, Elena Krugliakova, Georgia Sousouri, Sophia Snipes, Jelena Skorucak, Selina Schühle, Manuel Müller, Maria Laura Ferster, Giulia Da Poian, Walter Karlen, Reto Huber

**Affiliations:** Child Development Centre and Children’s Research Centre, University Children’s Hospital Zurich, University of Zurich, Zurich, Switzerland; Donders Institute for Brain, Cognition and Behaviour, Radboud University Medical Center, Nijmegen, Netherlands; Mobile Health Systems Lab, Department of Health Sciences and Technology, Institute of Robotics and Intelligent Systems, ETH Zurich, Zurich, Switzerland; Institute of Pharmacology & Toxicology, University of Zurich, Zurich, Switzerland; Sensory-Motor Systems Lab, Department of Health Sciences and Technology, ETH Zurich, Zurich, Switzerland; Institute of Biomedical Engineering, Faculty of Engineering, Computer Science and Psychology, Ulm University, Ulm, Germany; Department of Child and Adolescent Psychiatry and Psychotherapy, Psychiatric Hospital, University of Zurich, Zurich, Switzerland

**Keywords:** sleep, memory, auditory stimulation, closed-loop, phase-targeted, K-complex, acoustically evoked, memory consolidation, declarative memory, verbal memory, causality, evoked spindles, slow-wave-spindle coupling, high-density EEG

## Abstract

Over the past decade, phase-targeted auditory stimulation (PTAS), a neuromodulation approach which presents auditory stimuli locked to the ongoing phase of slow waves during sleep, has shown potential to enhance specific aspects of sleep functions. However, the complexity of PTAS responses complicates the establishment of causality between specific electroencephalographic events and observed benefits. Here, we used down-PTAS during sleep to specifically evoke the early, K-complex (KC)-like response following PTAS without leading to a sustained increase in slow-wave activity throughout the stimulation window. Over the course of two nights, one with down-PTAS, the other without, high-density electroencephalography (hd-EEG) was recorded from 14 young healthy adults. The early response exhibited striking similarities to evoked KCs and was associated with improved verbal memory consolidation via stimulus-evoked spindle events nested into the up-phase of ongoing 1 Hz waves in a central region. These findings suggest that the early, KC-like response is sufficient to boost memory, potentially by orchestrating aspects of the hippocampal-neocortical dialogue.

## Introduction

The phenomenon of sleep comprises characteristic neuronal states which seem critical for diverse physiological functions, including the re-normalization of synaptic weights [1], clearance of neurotoxic waste products that accumulate during wakefulness [2], and memory consolidation [3]. Surprisingly, despite decades of sleep research, the precise electrophysiological mechanisms that underlie these functions still remain a subject of ongoing debate.

When it comes to establishing causal relationships, neuromodulatory techniques, with their capacity to experimentally manipulate specific electroencephalographic events [4], offer a valuable means to dissect the precise contributions of electroencephalographic events to certain sleep functions. Presently, a substantial body of literature suggests that high-amplitude slow waves (0.5 – 4 Hz) and sleep spindles (11 – 16 Hz), the two most prominent electrophysiological oscillations in the electroencephalogram (EEG) during non-rapid-eye movement (NREM) sleep, are in some form linked to the aforementioned sleep functions [5–8]. However, particularly in human research, establishing direct, causal relationships between these events and any sleep function is challenging due to our limited understanding of the exact electrophysiological events induced by neuromodulatory techniques.

Consider phase-targeted auditory stimulation (PTAS) as an example — a straightforward and non-invasive technique that has received significant attention in the last decade [9]. The electrophysiological effects of PTAS appears to vary depending on the specific stimulation paradigm employed. For instance, when targeting the positive peak or ascending slope of slow waves (up-PTAS), it consistently results in an increase of slow-wave activity (SWA; spectral power between 0.5 and 4 Hz) or a re-distribution thereof towards time windows of stimulation [9–26]. In contrast, when targeting the negative peak or descending slope of slow waves (down-PTAS), the impact on SWA has yielded mixed results, sometimes leading to increases [24] and other times to decreases of SWA [9, 26, 27].

In addition, in a recent study conducted in our laboratory [16], we observed a dual-phase response when delivering stimuli in 6-second stimulation-opportunity (ON) windows, targeting the up-phase of slow waves. This dual-phase response encompassed an early and a late phase. The early phase commenced within the first two seconds following the presentation of the first stimulus of respective ON windows and was marked by a transient increase in SWA and sigma activity, closely resembling evoked K-complexes (KCs) [16]. Subsequently, the response evolved into a late phase, characterized by a sustained elevation in SWA persisting for an additional four seconds. Notably, these distinct responses were contingent on the sleep depth of the individual, with the late response primarily manifesting in individuals experiencing deeper sleep states. Consequently, from a mechanistic perspective, it becomes challenging to attribute behavioral outcomes observed after up-PTAS to one or the other response with certainty, as both can be concurrently in play, at least in deep sleepers. This is why the frequently documented enhancement in verbal declarative memory consolidation following up-PTAS [reviewed in 28, 29] actually offers limited insights into which manipulated aspect of slow-wave sleep contributed to this memory improvement.

Is there a way to dissect the early from the late response? When using up- and down-PTAS, at least conceptually, acoustic stimuli are played during periods of cortical firing or silence, respectively [30]. Given this distinction, it raises the possibility that the early and/or late response may also be tied to the targeted phase of slow waves. Therefore, in the presented work, we reanalyzed a previously published data set exploratively [31] to dissect the early and late response with respect to the targeted phase of slow waves. In this data set, participants underwent three separate laboratory nights: one with up-PTAS, another with down-PTAS, and a third night without any stimulation. We hypothesized that the early and/or late electrophysiological response would be contingent upon the targeted phase, implying distinct effects for up- and down-PTAS. Indeed, our analysis unveiled that the late response was a unique feature of up-PTAS and was absent following down-PTAS. In contrast, the early response was consistently observed after both up-PTAS and down-PTAS.

Next, in a new data set, we leveraged the specificity of down-PTAS towards the early response to study its effects on two specific domains of declarative memory (verbal and spatial), as well as alertness. By doing so, we can attribute any observed change in behaviour specifically to the early response following down-PTAS, a distinction which is confounded in studies employing up-PTAS. We hypothesized that down-PTAS, evoking solely the early response following PTAS, has no effect on memory and alertness. Our findings, however, revealed that the early response, along with enhanced cross-frequency coupling between slow waves and spindles, were associated with significant improvements in overnight verbal memory consolidation. Other cognitive domains tested were unaffected. These results provide direct, first experimental evidence supporting a causal relationship between the early response after PTAS, which closely resembled evoked KCs, and verbal declarative memory consolidation in healthy, young adults.

## Results

### Re-analysis of previously published data

#### Response profiles following up- and down-PTAS

In a first step, we tested whether the early and late response following PTAS depend on the targeted phase of slow waves. For this purpose, we re-analyzed high-density (hd-) EEG recordings (128 channels) of twelve participants, which were collected as part of a previous study (see methods). In short, participants underwent three experimental nights: two stimulation nights (STIM; one with up-PTAS, the other with down-PTAS), and a third night without stimulation (SHAM; all randomized and counterbalanced). In the SHAM night, stimulation flags, for both up- and down-PTAS, were saved without presenting actual tones. Stimuli (50 ms pink noise) were presented during NREM sleep (inter-stimulus interval (ISI) ≥0.5 s) in an ON|OFF window design: ON windows (6 s), allowing stimulation, took turns with OFF windows (6 s), withholding stimulation.

##### Auditory evoked response

We evaluated the phase-locked slow-wave response for both stimulation paradigms by computing the auditory evoked response (AER) time-locked to the first stimulus of a given ON window. Due to the locking of stimulation flags to a certain phase of slow waves, the AER was extracted by subtracting the mean waveform during SHAM (representing the mean targeted slow wave), from the mean waveform during STIM (reflecting a combined response of the mean targeted slow wave and the AER itself; **Fig. 1B**; refer to **Fig. 1A** to see the mean waveform during STIM and SHAM). Averaging the EEG of one frontal channel (Fz), the location where the largest AER can be expected, revealed a pronounced AER compared to SHAM after both up- (max. *g* = 1.97) and down-PTAS (max. *g* = 1.91). The AER was visually and statistically indistinguishable between stimulation paradigms, that is, between up- and down-PTAS (*p* ≥ .459 for all sample points, FDR corrected; max. *g* = 1.19), emphasizing the striking similarity between the early phase-locked response observed in both stimulation approaches.

**Fig. 1.**
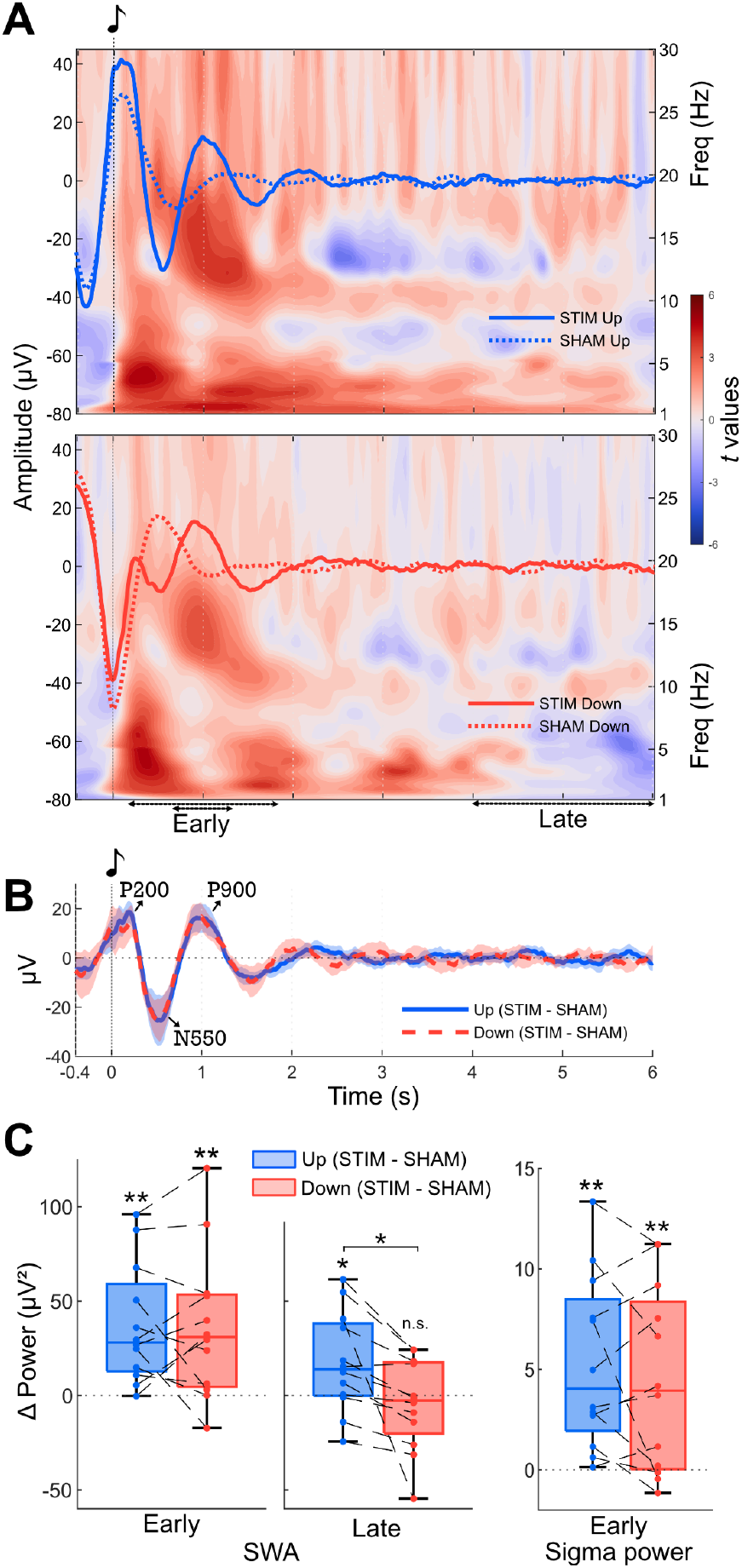
Comparison of early and late responses between up- and down-PTAS. **A)** Averaged EEG across stimuli (left y-axis) following actual (solid line) and SHAM stimulation (dotted line) in a night with up- (top) and down-PTAS (bottom). EEG data time-locked to the initial stimulus (0 s) of a given stimulation window (ON window) of channel Fz is displayed. The background illustrates the event-related spectral perturbation (ERSP; right y-axis) comparison with SHAM (paired *t* test). Arrows below the time axis signify time periods utilized for statistical comparisons in (C). The illustrated averaged waveforms and ERSP were computed using data from all windows regardless of the number of stimuli. **B)** Displayed is the auditory evoked response (AER, mean ± 95% CI), obtained by, for a given participant, subtracting the mean waveform following SHAM (representing the mean targeted slow wave), from the mean waveform during STIM (reflecting a combined response of the mean targeted slow wave and the AER itself). The AER was first computed for each participant individually and then averaged across all participants. The lines depict the average over participants. The P200, N550 and P900 are indicated with arrows. The illustrated AER was computed using data from all windows regardless of the number of stimuli. Of note, no considerable difference in the shape or amplitude of the AER was observed when considering windows with few or many stimuli only. **C)** Mean slow-wave activity (1 – 4 Hz; left) and sigma activity (12 – 16 Hz; right) averaged over all channels. Asterisks above box plots indicate statistically significant differences between STIM and SHAM (paired *t* test, * p < .05, ** p < .01, n.s. p ≥ .05). The early slow-wave (0.25 – 1.75 s) and sigma response (0.75 – 1.25 s) suggest a K-Complex-like response following both up- and down-PTAS. The later slow-wave response (4 – 6 s) is significantly stronger after up-PTAS compared to down-PTAS (as indicated by the horizontal bar).

##### Early slow-wave response

Slow-wave response profiles were further characterized by comparing slow-wave and sigma activity levels at different periods in time (**Fig. 1C**, refer to **Fig. 1A & B** to see the whole time-frequency spectrum). The analysis was performed separately for windows with few (1 – 4) and many (5 – 9) stimuli, as the number of stimuli relates directly to SWA levels present at times of stimulation. SWA levels [25] and sleep depth [16], in turn, have been shown to impact the resulting slow-wave response upon stimulation.

Compared to SHAM, early SWA averaged over all channels (1 – 4 Hz; STIM−SHAM; 0.25 – 1.75 s after stimulus onset) in windows with few stimuli was significantly larger following both up-PTAS, *t* (11) = 4.10, *p* = .002, *g* = 0.63, and down-PTAS, *t* (11) = 3.19, *p* = .009, *g* = 0.61 (**Fig. 1C**, middle panel). This early response was indistinguishable between stimulation paradigms, *t* (11) = 0.16, *p* = .877, *g* = 0.03.

Similarly, sigma activity levels (12 – 16 Hz; STIM−SHAM; 0.75 – 1.25 s after stimulus onset) were increased after both up-PTAS, *t* (11) = 4.30, *p* = .001, *g* = 1.04, and down-PTAS, *t* (11) = 3.33, *p* = .007, *g* = 0.85 (**Fig. 1C**, left panel). No statistical difference was observed between stimulation paradigms, *t* (11) = 0.99, *p* = .342, *g* = 0.18. Importantly, all findings related to the early slow-wave and sigma response remained consistent when considering all windows regardless of the number of stimuli or when using a restricted slow-wave range of 1 – 2 Hz instead of 1 – 4 Hz.

These results provide evidence that both stimulation paradigms evoke a similar early response. The presence of a subsequent sigma response, which followed the early slow-wave response, suggests a potential involvement of evoked KCs in driving the early response after PTAS [16, 32].

##### Late slow-wave response

Next, at the end of stimulation windows (4 – 6 s after stimulus onset), SWA was compared between stimulation paradigms in windows with numerous stimuli (**Fig. 1C**, right panel). When compared to SHAM, SWA was exclusively enhanced after up-PTAS, *t* (11) = 2.25, *p* = .046, *g* = 0.19, but not after down-PTAS, *t* (11) = −0.68, *p* = .509, *g* = −0.06. Furthermore, SWA (STIM−SHAM) after up-PTAS was significantly larger compared to down-PTAS, *t* (11) = 2.99, *p* = .012, *g* = 0.81.

To summarize, our findings highlight the significance of the targeted phase of slow waves in relation to the late, but not to the early response following PTAS. Both types of PTAS can elicit a robust early response, potentially driven by evoked KCs. However, the sustained enhancement of SWA occurring later in the stimulation window appears to be exclusive to up-PTAS.

### Analyses of new data set

#### Effects of down-PTAS on EEG and memory

Building on these findings, our objectives were twofold. By applying down-PTAS in a new sample, we set out to 1) gain a deeper understanding of the EEG characteristics of the early response following down-PTAS and 2) examine its potential link to declarative memory consolidation and alertness. Hd-EEG recordings from 14 participants were analyzed. Participants underwent two experimental nights, one with down-PTAS (STIM), the other without stimulation (SHAM; randomized and counterbalanced). Stimuli (50 ms pink noise) were presented during NREM sleep (ISI ≥0.5 s) in 16 s ON and 8 s OFF windows, following the signal of a target electrode next to C3 (see methods). The median ISI of all stimuli ranged between 0.96 and 1.41 seconds within participants (mean of 1.27 seconds across participants).

#### EEG characteristics of the early response

The early response following the first stimulus of ON windows is characterized by a guaranteed stimulation break prior to the first stimulus (OFF window). Capitalizing on the extended duration of the ON windows, we conducted the following analyses with reference to isolated stimuli, ensuring at least 5 seconds without stimulation preceding each stimulus to allow adequate time for the refractory period of KCs to subside [33, 34]. Across participants, 184 to 859 stimuli met this criterion (mean±sd = 502.75±161.22 stimuli), representing an increase of 5.74 to 26.27% (mean±sd = 17.12±5.18%) compared to the number of first stimuli.

Having observed initial indications for the KC-like nature of the early response following PTAS in our re-analysis of the previous data set (**Fig. 1)**, we sought additional evidence to support this resemblance in the new data set. For this purpose, we first evaluated the AER, specifically focusing on the N550 component, which is deemed essential for classifying a response as KC-like [35].

##### Auditory evoked potential and N550

Averaging the EEG of one frontal channel (Fz) time-locked to isolated stimuli demonstrated a prominent AER compared to SHAM (*p* < .05, FDR corrected; max. *g* = 2.36; **Fig. 2A**). The AER exhibited all classical sleep AER components (P200, N350, N550 and P900). When identifying the N550 component in channel Fz and evaluating its amplitude across channels, it was found to be largest over fronto-central regions (*p* < .05, cluster corrected; max. *g* = 2.27; **Fig. 2B**), matching the anticipated topography of the N550 component of KCs [36, 37].

**Fig. 2.**
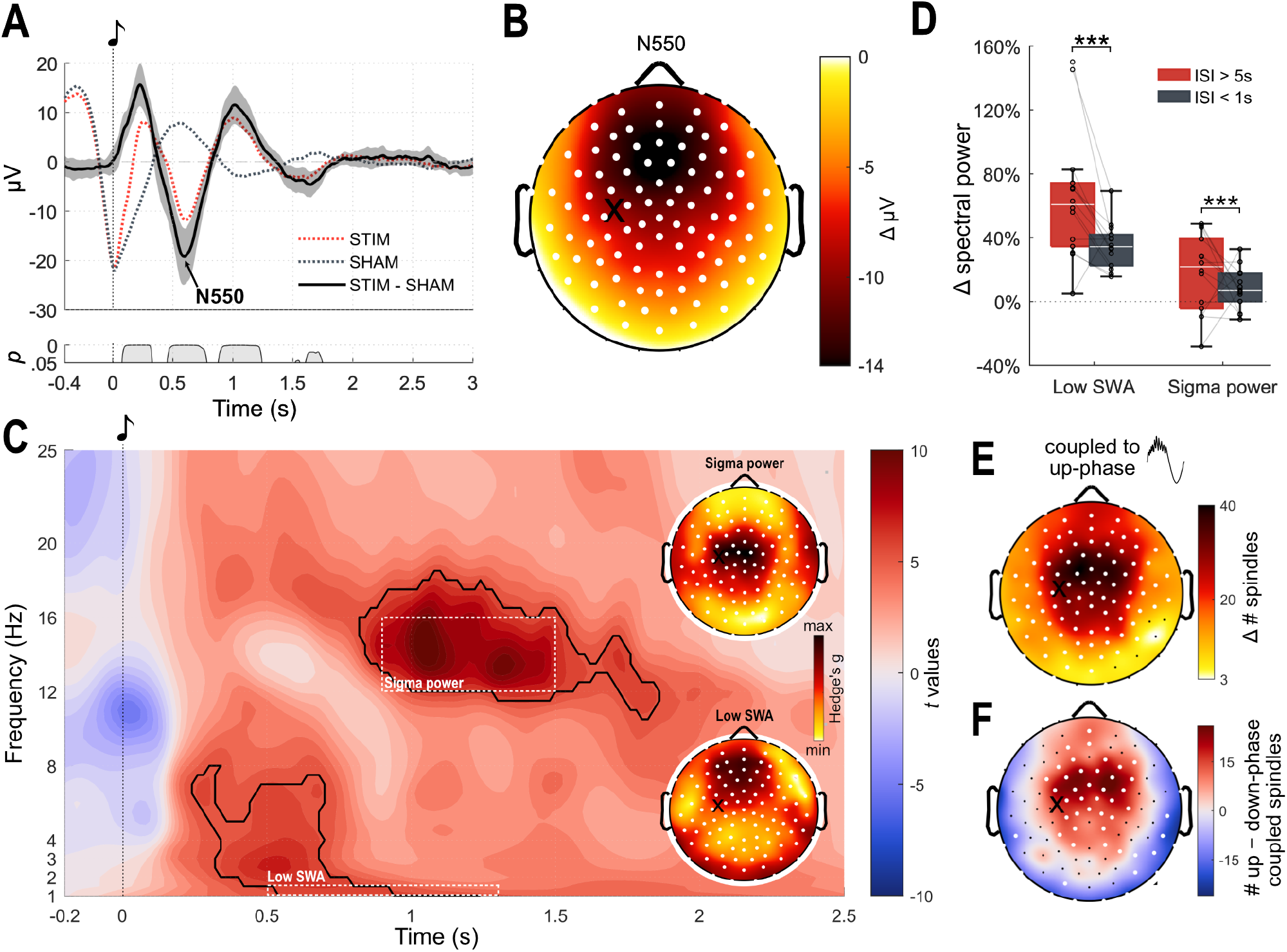
Indications for a K-complex-like response. In all panels, time 0 s corresponds to the onset of isolated stimuli, that is, stimuli with an inter-stimulus interval (ISI) ≥ 5 s. **A)** Averaged EEG across stimuli (dotted lines) and auditory evoked response (AER, mean ± 95% CI, thick black line) of channel Fz as in **Fig. 1**. The bottom panel displays significant differences between conditions (paired *t* test, FDR corrected). **B)** Topography of the difference in N550 amplitude between conditions (STIM −SHAM). The latency of the N550 component was determined individually for each night, based on the averaged EEG of the channel average in a predefined frontal region of interest (ROI, see **Supplementary Fig. 8)**. White dots represent electrodes with significant differences between conditions (paired *t* test, cluster corrected, see methods); the black cross indicates the target electrode. **C)** Normalized (participant- and frequency-wise, by the trial average of the entire time window) event-related spectral perturbation (ERSP). The ERSP of the channel average in a predefined frontal ROI (see **Supplementary Fig. 8)** is shown. Contour lines signify significant differences between conditions (paired *t* test, cluster corrected, see methods). The topography of effect sizes (Hedge’s g) of the mean sigma (900 – 1500 ms; 12 – 16 Hz; Hedge’s g value range: 0.46−1.21) and low slow-wave response (500 – 1300 ms; 1 – 1.5 Hz; Hedge’s g value range: 0.93−2.67) is depicted at the top and bottom, respectively. White dots and black cross can be interpreted as in (B). **D)** Low SWA and sigma power (as in C), averaged over channels, after isolated (ISI≥5 s; in red) and consecutive stimuli (ISI < 1 s; in gray). Circles represent values of single participants. Stars indicate significant differences (paired *t* test, *** p < .001). **E)** Topography depicting the difference in the number of evoked spindles between conditions (STIM −SHAM) detected 2 s following isolated stimuli coupling to the positive half-wave (up-phase) of 1 Hz waves (paired *t* test, cluster corrected, see methods). White dots and black cross can be interpreted as in (B). **F)** Topography displaying the difference in the number of up- and down-phase coupled spindles (up −down), contrasted between conditions (STIM −SHAM; paired *t* test, cluster corrected, see methods). White dots and black cross can be interpreted as in (B).

##### Event-related spectral perturbation

Alongside the close relationship between KCs and sleep spindles, it has been noted that evoked KCs exhibit a distinctive spectral profile, featuring increases in the theta, alpha, sigma, and beta bands [32, 38, 39].

To evaluate this spectral profile, the event-related spectral perturbation (ERSP) was computed time-locked to isolated stimuli within a predefined frontal region of interest (ROI, **Supplementary Fig. 8)**. The analysis revealed a significant increase in sigma activity (12 – 16 Hz) around 0.9 – 1.5 s after stimulus presentation in comparison to SHAM (*p* < .001, cluster corrected, 99.99th percentile; **Fig. 2C**). Additionally, the stimuli evoked an early response in the low slow-wave band (1 – 1.5 Hz) around 0.5 – 1.3 s, as well as in the high slow-wave (2 – 4 Hz) and theta band (4 – 8 Hz) around 0.3 – 0.7 s after stimulus presentation. Further topographic analyses of the low slow-wave and sigma response revealed that the increase in SWA and sigma activity was global (*p* < .05, cluster corrected), yet varied topographically in terms of effect sizes (topographies in **Fig. 2C**). The increase in low SWA was most pronounced over fronto-central regions (max. *g* = 2.62), while the increase in sigma activity was greatest over central areas (max. *g* = 1.21), aligning well with the expected topography of the slow-wave and spindle components of the evoked response.

##### Response after subsequent stimuli

Since KCs exhibit a refractory period [40], a larger increase in spectral power, if attributable to evoked KCs, would be anticipated to occur after isolated stimuli compared to consecutive stimuli, here defined by having another tone the second preceding the stimulus. Across participants, 396 to 1896 consecutive stimuli meeting this criterion were identified (mean±sd = 883.39±349.81 stimuli). To examine whether isolated stimuli induced a larger increase in spectral power, the increase in low SWA and sigma activity in response to isolated and consecutive stimuli was compared in a predefined frontal ROI. Both the slow-wave, *t* (13) = 5.45, *p* < .001, *g* = 1.32, and sigma response, *t* (13) = 6.48, *p* < .001, *g* = 1.90, were stronger after isolated compared to consecutive stimuli (**Fig. 2D**). As the first stimulus in an ON window is, per definition, an isolated stimulus, this distinction is also evident when comparing the spectral response following the first as opposed to the second stimulus within an ON window. Specifically, SWA, *t* (13) = 3.68, *p* = .003, *g* = 0.57, and sigma activity, *t* (13) = 4.11, *p* = .002, *g* = 1.04, were higher after the first compared to the second stimulus (data not shown). Similarly, the amplitude of the N550 component in channel Fz was stronger after isolated compared to consecutive stimuli, *t* (13) = 5.85, *p* < .001, *g* = 1.60 (data not shown). Collectively, these findings suggest that stimuli occurring following longer stimulation breaks elicit a more pronounced KC-like response.

##### Stimulus-induced spindle events

The question arises whether the observed enhancement in sigma activity following isolated stimuli (**Fig. 2C**) directly translates into stimulus-evoked spindle events. Since KCs are often precursors to sleep spindles [41], the presence of stimulus-evoked spindles would further provide evidence supporting the KC-like nature of the evoked response.

Discrete spindle events occurring within 2 seconds following isolated stimuli were detected using a previously described method [42]. Compared to SHAM, a global increase in the number of detected spindles was observed after stimulation, mirroring the same topographical distribution as depicted in **Fig. 2E** (*p* < .05, cluster corrected; max. increase of 55.71 spindles; max. *g* = 0.94; data not shown). This effect persisted when controlling for the number of delivered stimuli (max. *g* = 1.43) or the total number of spindles detected during the entire night (max. *g* = 1.17).

##### Coupling of stimulus-induced spindle events

Given the observed enhancement in sigma activity occurring 0.9 – 1.5 s following stimulus presentation (**Fig. 2C**), coinciding with the slow P900 component of the AER (**Fig. 2A**), evoked spindle events may exhibit preferential coupling to the up-phase of underlying slow waves. To test this, the underlying phase of 1 Hz waves was extracted at the point of maximum amplitude of detected spindles.

Compared to SHAM, a global increase in the number of spindles nesting within the up-phase of 1 Hz waves was observed (*p* < .05, cluster corrected; max. difference of 39.64 spindles; max. *g* = 1.04; **Fig. 2E**). Notably, significantly more spindles coupled to the up-compared to down-phase of 1 Hz waves in a frontal cluster of 35 electrodes (*p* < .05, cluster corrected; max. difference of 27.00 spindles; max. *g* = 0.87), while a peripheral cluster of 17 electrodes showed a reduction (*p* < .05, cluster corrected; min. difference of −29 spindles; min. *g* = −1.15; **Fig. 2F**). Relative to the total number of evoked spindles, the proportion of up-phase-coupled spindles was higher in a cluster of 8 frontal electrodes, yet this effect did not survive cluster correction (*p* < .05, max. difference of 8.55%; max. *g* = 1.08; data not shown). These findings collectively suggest that the previously observed increase in sigma power in the vicinity of the slow P900 component of the AER does indeed materialize as discrete spindle events which preferentially occur fronto-centrally within the up-phase of underlying slow waves.

#### Spectral power changes throughout and beyond the stimulation window

The preceding analyses indicated that the early response following PTAS indeed resembles a KC-like response. However, the effects of stimuli later in the stimulation window have not been fully explored. Hence, in a next step, we analyzed the EEG response throughout the entire ON|OFF window, thereby focusing on changes in SWA.

##### SWA response within stimulation windows

The 16 s ON and 8 s OFF windows were divided into equally long windows of 8 s duration in order to compare SWA (1 – 4.5 Hz) between conditions (STIM−SHAM). Consistent with the previous findings of a KC-like response to the first and isolated stimuli, SWA was globally increased at the beginning of stimulation windows (ON1; *p* < .05, cluster corrected; data not shown) but not later on (ON2 & OFF; all *p* < .05). This makes sense given that, across participants, 60.68 to 88.00% (mean±sd = 73.16±6.30%) of isolated stimuli occurred within the first 6 s of ON windows. To investigate a potential dose-response relationship and account for ongoing SWA during stimulation, we further divided ON|OFF-window pairs based on the number of stimuli presented within the first 6 s of stimulation. Intriguingly, the observed increase in SWA during ON1 was most pronounced in stimulation windows with 1 – 2 stimuli (max. *g* = 3.75), followed by those with 3 – 4 stimuli (max. *g* = 2.04), and was completely absent in windows with 5 or more stimuli (*p* < .05, cluster corrected; max. *g* = 1.27; **Fig. 3A**). These results, while initially surprising, can be explained by the elevated arousal threshold during deeper sleep stages [43]. It is during lighter sleep stages that soft stimuli are more likely to elicit KCs. Additionally, the absence of SWA enhancements during windows with numerous stimuli supports the concept that down-PTAS, as opposed to up-PTAS, lacks the capacity to effectively drive endogenous SWA apart from KC induction.

**Fig. 3.**
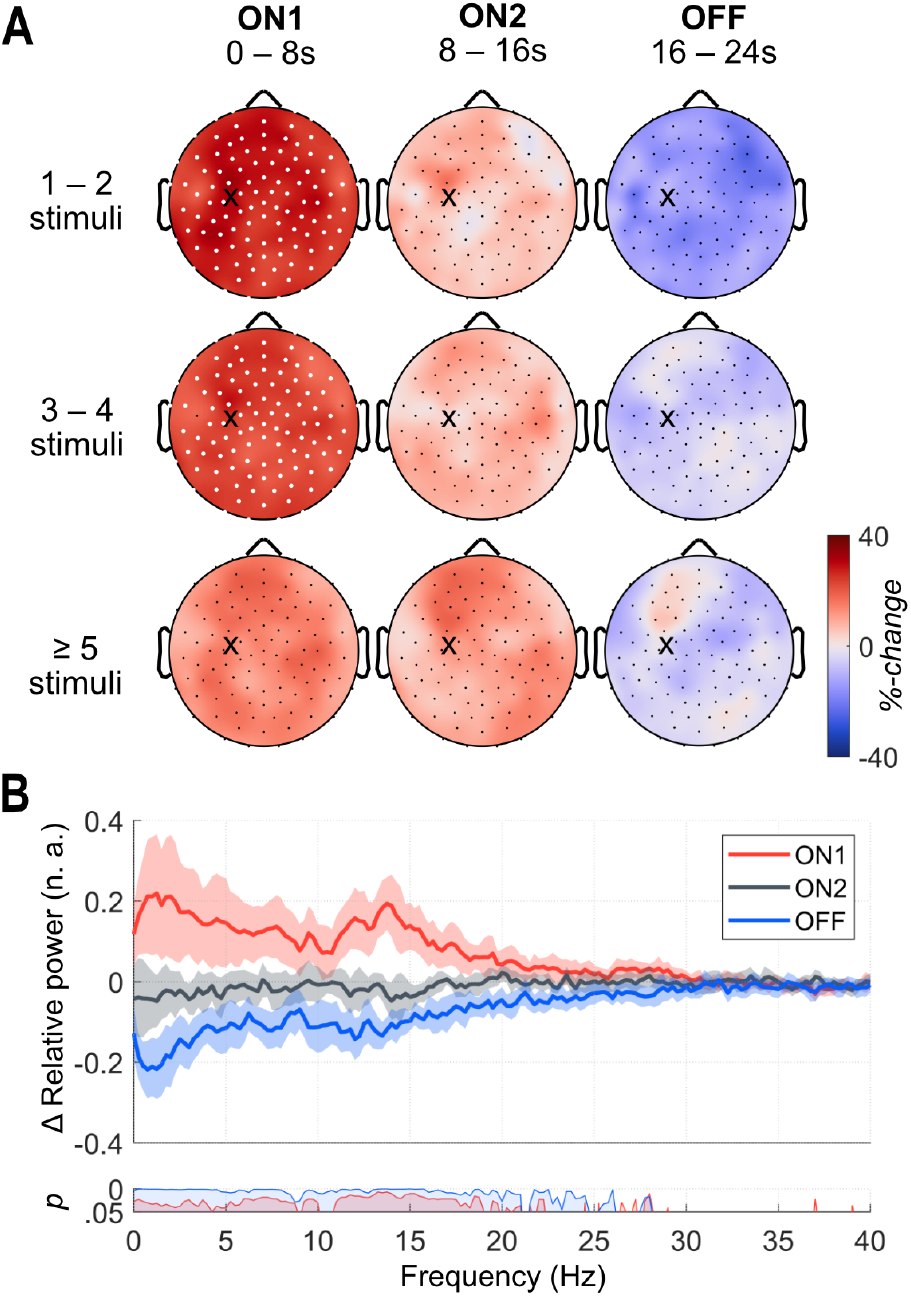
Spectral power in ON and OFF windows. **A)** Topography of slow-wave activity (SWA; 1 – 4.5 Hz) within 8 s window snippets (ON1, ON2, OFF) containing few (1 – 2), medium (3 – 4) and many (≥5) stimuli within the first 6 s in ON1. The difference in SWA (STIM−SHAM) in relation to the average of both nights is displayed. White dots represent electrodes with significant differences between conditions (paired *t* test, cluster corrected, see methods); the black cross indicates the target electrode. **B)** Relative power spectral density (PSD, mean ± 95%CI) of 8 s window snippets (ON1, ON2, OFF) belonging to ON|OFF-window pairs which had 1 – 2 stimuli within the first 6 s in ON1. PSD values within window snippets are normalized to the channel average of the entire ON|OFF window (window- and frequency-wise). Relative PSD values thus indicate a redistribution of PSD within windows. The average PSD of all electrodes is depicted. Bottom panel displays significant differences between conditions (paired *t* test, FDR corrected).

The limited duration of SWA enhancements already suggests the temporal specificity of the stimulation effect. Comparing the time spent in different sleep stages across condition (STIM−SHAM) confirmed the absence of any significant alterations in sleep architecture (paired *t* test, FDR corrected, all *p* ≥ .05; **Supplementary Fig. 3D**). Additionally, no difference was observed in overnight SWA, as indicated by mean SWA across all NREM epochs (paired *t* test, cluster corrected, all *p* ≥ .05; **Supplementary Fig. 3A** − **C**).

##### Re-distribution of spectral power within ON|OFF window pairs

To investigate whether the increase in spectral power at the beginning of stimulation windows was specific to SWA, we assessed the redistribution of spectral power across the 0 – 40 Hz frequency range. For a given frequency, PSD values within ON1, ON2, and OFF windows were normalized by the channel average PSD of the respective entire ON|OFF window and compared between conditions (STIM−SHAM). Most frequencies up to 18 Hz, with the exception of alpha activity (∼10 Hz), were redistributed towards ON1 and away from OFF (**Fig. 3B**). These findings suggest that the early response following PTAS not only enhances SWA compared to SHAM, but also causes a shift in SWA, theta, and sigma activity towards the beginning of stimulation windows, where evoked KCs would be concentrated.

#### Consequences on memory and alertness

Although down-PTAS induced a prominent early EEG response resembling a KC, its functional significance remains unclear. To further investigate this, we examined the effect of down-PTAS on verbal and spatial declarative memory consolidation, as well as alertness.

##### Memory

Memory was assessed using an associated word-pair memory task (40 word-pairs; **Supplementary Fig. 6**) and an object-location task (15 object-locations; **Supplementary Fig. 7**). The difference in recalled items before and after sleep was calculated as the score for overnight memory consolidation. Additionally, participants learned and directly recalled a new list of 40 word pairs in the morning to asses their encoding capabilities after down-PTAS (illustration of procedure in **Supplementary Fig. 5D & E**). For the associated word-pair task, auditory stimulation significantly improved overnight verbal memory consolidation, *t* (12) = 2.58, *p* = .023, *d* = 0.72 (**Fig. 4A**). No significant differences were found for encoding capabilities of new word-pairs, *t* (12) = 0.52, *p* = .606, *d* = 0.15 (**Supplementary Fig. 5A**), or for the consolidation of object-locations (**Supplementary Fig. 5B**), *t* (11) = 0.00, *p* = 1.000, *d* = 0.00.

**Fig. 4.**
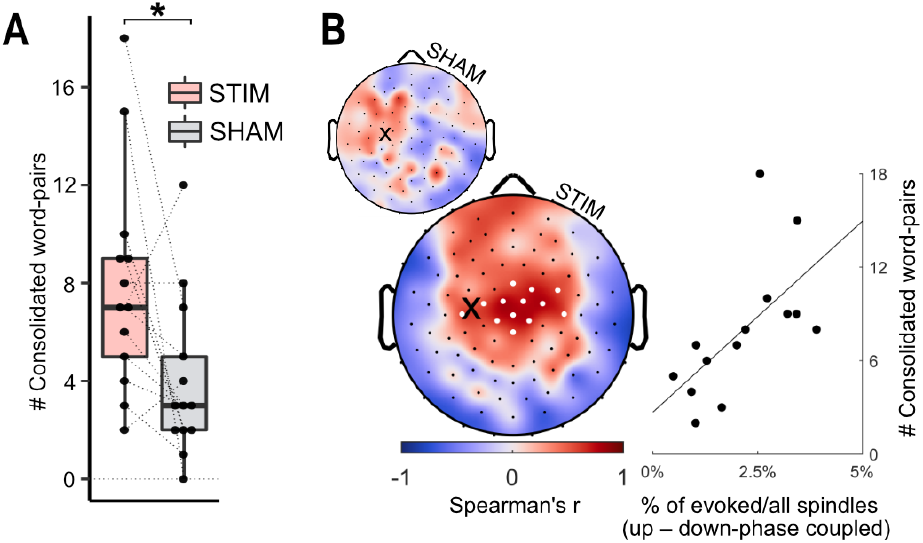
Verbal memory consolidation. **A)** Overnight memory improvement in the associated word-pair memory task (morning − evening) after a night with (STIM) and without auditory stimulation (SHAM). Dots represent the performance of single participants. **B)** (Left) Detected spindles were classified based on their coupling to the positive (up-phase) or negative (down-phase) halfwave of 1 Hz waves. Depicted is the correlation between the number of consolidated word-pairs and the difference in the number of up- and down-phase coupled spindles, expressed relative to the total number of spindles detected during the entire night, separate for SHAM (top) and STIM (bottom). White dots represent electrodes with significant correlations (p < .05; cluster corrected, see methods); the black cross indicates the target electrode. (Right) Spearman’s rank correlation coefficient, averaged over significant electrodes (for STIM).

##### Alertness

To investigate the impact of down-PTAS on alertness, participants underwent a simple reaction time task before and after sleep. Both phasic and tonic alertness were assessed (see methods). The standard deviation in reaction times measured in the morning was subtracted from those measured in the evening to assess alertness. Our results indicate that down-PTAS had no significant effect on either phasic, *t* (12) = −0.23, *p* = .823, *d* = −0.06, or tonic alertness, *t* (12) = −0.24, *p* = .814, *d* = −0.07 (**Supplementary Fig. 5C**). These results remain consistent when evaluating alertness based on the average reaction times instead of the standard deviation.

#### Positive relationship between detected sleep spindles and memory performance

Given the growing body of evidence supporting the involvement of sleep spindles in sleep-dependent memory processes, we aimed to investigate the relationship between stimulus-evoked spindles and verbal declarative memory performance following down-PTAS using Spearman’s rank correlation coefficient.

The absolute number of stimulus-evoked sleep spindles did not significantly correlate with verbal memory performance (all *p* > .05, cluster corrected; mean *r*(12) = .38 across all channels; **Supplementary Fig. 1A**). However, normalizing the absolute number of stimulus-evoked spindles to the total number of overnight spindles revealed a positive correlation between the proportion of stimulus-evoked spindles and the consolidation of verbal memory, exclusively in the stimulation night, evident in a global cluster of 99 channels (*p* < .05, cluster corrected; mean *r*(12) = .68, mean *p* = .010; **Supplementary Fig. 1A**).

##### The relevance of slow-wave-spindle coupling

Previous studies have specifically linked sleep spindles that couple to the up-phase of slow waves to enhanced memory consolidation [44]. Our data, however, revealed a positive relationship to verbal memory for both up-phase (*p* < .05, cluster corrected; mean *r*(12) = .68, mean *p* = .010 in a cluster of 71 channels; **Supplementary Fig. 1B**) and down-phase coupled spindles (*p* < .05, cluster corrected; mean *r*(12) = .66, mean *p* = .015 in a cluster of 77 channels; **Supplementary Fig. 1B**). This indicates that stimulus-evoked sleep spindles enhance verbal memory consolidation regardless of their phase coupling.

Nevertheless the distinct impact of up-phase coupling becomes evident when directly comparing the normalized number of up-phase to down-phase coupled spindles. A greater proportion of up-phase compared to down-phase coupled spindles correlated significantly with enhanced verbal memory performance within a central cluster of 13 channels, observed exclusively during the stimulation night (*p* < .05, cluster corrected; mean *r*(12) = .66, mean *p* = .019, **Fig. 4B**).

Consistent with studies demonstrating stronger coupling to slow waves at around 1 Hz [e.g., 45], this positive correlation was evident solely for sleep spindles coupled to 1 Hz slow waves, moved to a frontal location for spindles coupled to 2 Hz slow waves, observed in a cluster of 6 channels (*p* < .05, cluster corrected; mean *r*(12) = .64, mean *p* = .015; **Supplementary Fig. 1C**), and was weakest when considering the phase of faster slow waves (3 – 4 Hz; **Supplementary Fig. 1C**).

These findings collectively suggest that while stimulus-evoked spindles appear to generally promote verbal memory consolidation, those coupled to the up-phase of 1 Hz waves may be particularly beneficial. This positive relationship provides a potential mechanism for how the early KC-like response following PTAS interacts with declarative memory consolidation.

## Discussion

Our study provides compelling evidence supporting a causal relationship between the early EEG response following PTAS, resembling evoked KCs, and the consolidation of verbal declarative memory, as evidenced by a significant improvement in verbal memory performance among healthy adults. Opting for down-over up-PTAS, we effectively evoked the early EEG response following PTAS while concurrently eliminating possible interference from later EEG responses as typically observed after up-PTAS.

This approach allowed for a direct examination of the causal impact exerted by specifically the early EEG response on memory and alertness.

Our results demonstrate that down-PTAS can successfully improve the consolidation of verbal declarative memory, while spatial declarative memory and alertness were not affected. In the EEG, we observed a robust early response which exhibited striking resemblance to evoked KCs and was accompanied by evoked spindle events that preferentially coupled to the up-phase of slow waves. Importantly, the proportion of acoustically evoked spindles relative to the total number of spindles during the entire night, particularly when coupled to the up-phase of 1 Hz waves, was positively correlated with the consolidation for previously learned word-pairs.

### The missing link between auditory stimulation and improved memory consolidation

While numerous studies could demonstrate memory enhancements after auditory stimulation [reviewed in 28, 29], they have often fallen short in disentangling which specific electrophysiological response following stimulation is actually responsible for the improvements in memory. In addressing this critical gap, our study makes a significant contribution by

1. implementing a stimulation approach, i.e., down-PTAS, that effectively isolates the early response while minimizing confounding late responses **Fig. 1**,
2. conducting a comprehensive investigation of the relationships between EEG responses following stimulation and the observed improvements in memory consolidation.

To the best of our knowledge, this study represents the first direct experimental evidence supporting the specific causal role of the early, KC-like response following PTAS in verbal memory consolidation in healthy adults.

### The early response does not universally improve all aspects of sleep function

Given the rather global enhancement of SWA and sigma activity during the early response following PTAS, its exclusive positive impact on verbal declarative memory, without extending to visuo-spatial memory consolidation, verbal encoding capabilities, or alertness, is rather surprising, albeit generally consistent with the literature on up-PTAS [22].

Notably, our study identified a significant correlation between verbal memory performance and the proportion of evoked spindles relative to the total spindle count during the entire night, rather than the absolute number of evoked spindles. This finding is particularly intriguing given the complex relationship between sleep spindle characteristics and verbal declarative memory [46], with both positive [47–50] and negative [51] associations being reported depending on the specific spindle feature under investigation. Consistent with the notion that distinct sleep spindles, such as those differentiated based on frequency, fulfill distinct functional roles [46, 52], it appears that those that are evoked through auditory stimulation generally promote verbal but not visuo-spatial memory consolidation.

Exploring the precise reason why the early response favored verbal memory consolidation over visuo-spatial memory remains a subject of speculation at this stage, yet aligns with the prevailing concept that the mechanisms governing the consolidation of distinct memory domains vary [53]. Potentially, the relation of the memory trace domain (verbal) to the domain of the stimulus (auditory) may be of importance. Generally, especially in the context of auditory stimulation as a sleep-enhancing tool, our results emphasize the importance of considering both the prevailing brain state during stimulation [16] and the specific set of stimulation parameters (including, but not limited to, the targeted phase of slow waves), when estimating electrophysiological response(s) following PTAS and its functional consequences.

### Down-PTAS does not inevitably improve memory

Interestingly, the only other study which investigated the effect of down-PTAS on memory consolidation reported no improvements in either memory consolidation or SWA levels following down-PTAS [9]. A plausible explanation for this discrepancy lies in the difference in the stimulation protocol used between the two studies. Ngo et al. 9 observed an enhancement in SWA and memory recall when using up-PTAS with a two-click design and an ISI of 1.075 seconds. Conversely, when down-PTAS was employed with a reduced ISI of 0.55 seconds, the enhancements in SWA and memory recall were absent. Evidently, the targeted phase and the selection of the ISI are inherently intertwined, posing limitations in isolating their individual contributions. Can the change in ISI explain the absence of memory benefits after down-PTAS?

It is well-documented that sleep spindles reduce sensory responsiveness towards external stimuli [54]. Importantly, in our study, we observed a robust spindle response 0.9 – 1.5 s following stimulus presentation. In the study of Ngo et al. 9, exclusively during up-but not during down-PTAS the second click temporally aligns with the evoked spindle response, which could mediate the potentially arousing effect of a second click. In our study, in which down-PTAS improved memory performance, the median ISI ranged between 0.96 and 1.41 seconds within participants (mean of 1.27 seconds across participants), aligning well with the spindle response.

Hence, while all stimulation protocols likely evoked KCs, the relationship between KCs and arousals appears to be of importance in their functional role towards memory consolidation. This observation also aligns with the notion that KCs seem to have a dual functional role, as highlighted in the continuous discussion regarding their contribution to both arousal and sleep protection [55].

### A possible mechanism how auditory stimulation facilitates memory consolidation

Rather than increasing overall SWA or altering sleep architecture, the temporal scope of the early, KC-like response was distinctly confined to the period of stimulation. During that confined time period, the early response following PTAS contributed to the precise orchestration of the timing between slow waves and sleep spindles, evidenced as stimulus-evoked spindles preferentially nesting within the up-phase of slow waves. Importantly the proportion of stimulus-evoked up-phase coupled spindles relative to the proportion of down-phase coupled spindles exhibited a positive correlation to verbal memory consolidation within a central cluster specifically. These results provide a possible mechanism how the early, KC-like response may have facilitated verbal memory consolidation. According to prevailing theories on memory consolidation, such as the hippocampal memory indexing theory [56] or the current active system consolidation concept [5], it is the synchronization between slow waves, sleep spindles, and hippocampal ripples, along with their temporal hierarchy, that reflect the hippocampal-neocortical dialogue crucial for memory consolidation [reviewed in 57]. Consequently, our data supports the conceptual framework that the early, KC-like response may actively participate in the hippocampal-neocortical dialogue, thereby strengthening neuronal connections underlying the memory trace [reviewed in 58].

## Conclusions

By experimentally inducing specifically the early EEG response following PTAS and simultaneously observing improved verbal declarative memory consolidation in young healthy adults, this study provides compelling and direct evidence establishing a causal relationship between this response, strongly resembling evoked KCs, and the consolidation of verbal declarative memory. Importantly, in contrast to studies which employed up-PTAS, the use of down-PTAS effectively minimized the interference of later responses observed after up-PTAS, which could otherwise have benefited sleep functions as well. Stimulus-evoked spindles nesting within the up-phase of the underlying slow wave present as a plausible mechanism through which the early response enhances memory consolidation, aligning well with prevailing theories on memory consolidation during sleep.

## Limitations

This study encompasses certain limitations that warrant acknowledgment. One of the primary limitations is the rather small number of participants, a consequence of the extensive effort required to conduct overnight sleep EEG experiments in a laboratory setting. This limitation may restrict the generalizability of our results and could potentially impact the statistical power of our analyses. Additionally, our decision to use comfortable over-ear headphones, while intended to enhance participant comfort, introduced variable pressure on the ears depending on the sleeping positions of the participants. This variability likely affected the perceived volume of the auditory stimuli, adding another layer of complexity to our findings. Finally, while sleep spindles were detected using a well-established procedure and the outcome of single channels was manually verified, a more liberal threshold was necessary to ensure an adequate number spindle events following the restricted time window following isolated stimuli. This decision may have affected the reliability of spindle event detection.

## Future directions

Optimizing stimulus delivery to evoke KCs could be further enhanced in future stimulation protocols by incorporating long and irregular ISIs, integrating deviant stimuli, and considering the brain state, for example, whether tones are played during the ascending or descending slope of a sleep cycle [59].

## ^2^Abbreviations

(KC): K-complex
(SWA): slow-wave activity
(PTAS): phase-targeted auditory stimulation
(hd-EEG): high-density electroencephalography
(EMG): electromyography
(ecg): electrocardiography
(NREM): non-rapid eye movement
(REM): rapid eye movement
(ISI): inter-stimulus interval
(FDR): false discovery rate
(AER): auditory evoked response
(ERSP): event-related spectral potential
(ROI): region of interest
(AASM): American Academy of Sleep Medicine
(PSD): power spectral density
(FIR): finite impulse response
(M): mean
(SD): standard deviation
(g): Hedge’s g
(d): Cohen’s d

## Acknowledgments

We extend our gratitude to every member of Prof. Reto Huber’s lab and the entire SleepLoop consortium for their constructive feedback and insightful discussions. Special thanks are due to Maria E. Dimitriades for her valuable discussions and meticulous proofreading. We further express our sincere gratitude to Simone Accascina for providing invaluable technical assistance for our custom recruiting website. We express our appreciation to Cécile Abati, Tamara Weil, Paulina Schmiedeberg, Nicole Meier, Vanessa Kasties, Veronika Voskresenska, and Gila Norup for their valuable contributions to data collection. Lastly, we are grateful to all the study participants for their participation and cooperation. This work was supported by the Swiss National Science Foundation (SNF) under grant number 320030_179443 and is part of the HMZ Flagship grant «SleepLoop» under the umbrella of «Hochschulmedizin Zürich», Switzerland.

## Author contributions

Author contributions follow the Contributor Roles Taxonomy. **Conceptualization**: SL and RH; **Data curation**: SL, SSch, and MM (new data); EK, JS, SL, SS (existing data) **Formal analysis**: SL (new & existing data) and EK (existing data); **Funding aquisition**: RH; **Investigation**: SL, SSch, and MM (new data); EK, JS, SL, and SS (existing data); **Project administration**: SL and RH; **Resources**: RH; **Software**: MLF, GP, GS, WK, SS, SL, and MM; **Supervision**: RH; **Visualization**: SL; **Writing**: SL (original draft); all authors (review and editing).

## Data availability

The data sets used and/or analysed during the current study available from the corresponding author on reasonable request.

## Methods

### Re-analysis of existing data

To investigate the hypothesis that up- and down-PTAS lead to distinct profiles of slow-wave responses, we conducted a re-analysis of hd-EEG recordings from a subset of 12 out of 18 participants aged between 18.38 and 26.69 years (mean±sd = 23.83±1.51 years; 6 male, 6 female). The 12 participants had previously undergone the study procedure comprising three experimental nights: one with up-PTAS, one with down-PTAS, and one without stimulation (SHAM) [31]. The data collected after a night with up-PTAS [17, 31] and down-PTAS [31] had been previously analyzed for different research purposes.

#### EEG analyses

EEG analyses of existing data were performed in Matlab (R2017b & R2023a; The Math-Works, Inc., Natick, Massachusetts). The auditory evoked potential was computed as described later. Time-frequency analyses were performed using Hanning-window-tapered wavelets and sliding time windows with a time step of 0.02 seconds. The length of the sliding window was adjusted to ensure that it would fit 4 full cycles for frequencies between 0.75 and 5 Hz, and 6 full cycles for frequencies between 5 and 30 Hz. The average-referenced EEG data was aligned to the onset of the first stimulus within an ON window and used as input for Fieldtrip’s *ft_freqanalysis()* function with the *mtmconvol* option. ON windows were categorized based on the number of stimuli occurring within each window, either between 1 and 4 stimuli or between 5 and 9 stimuli. The average ERSP over ON windows was computed and stored for further analysis.

### Collection of new data

New data was collected employing down-PTAS to investigate the role of the early response following PTAS on two domains of declarative memory consolidation. Additionally, the early response was investigated in relation to alertness.

### Participants

A total of 17 participants were initially recruited, with 14 (4 male, 10 female) successfully completing the study. All participants were right-handed and aged between 18.38 and 26.69 years (mean±sd = 23.25±2.53 years). Three participants dropped out due to reasons including disregarding bedtimes (N = 1), poor sleep efficiency (<70%, N = 1), and technical issues (N = 1).

All participants except for one were native Swiss-German speakers. Prior to the study, in an anonymous online-screening, all participants self-disclosed their medication-free status, low consumption of caffeine and alcohol, and abstinence from nicotine and other drugs. They also reported no personal or family history of neurological or psychiatric disorders, including sleep disorders. Participants affirmed themselves as good sleepers with an average chronotype and were excluded from the study if they had engaged in shift work, daytime napping, or traveled across more than two time zones within the previous 30 days.

The study was approved by the local ethics committee (Kantonale Ethikkommission Zürich, KEK-ZH, BASEC 2019-02134) and was conducted in adherence to the principles outlined in the Declaration of Helsinki. Written informed consent was obtained from all participants before their participation in the study. Data collection took place between August 2020 and April 2021.

### Study design and procedure

Eligible participants initially followed a predetermined sleep schedule that matched their laboratory bedtimes for ∼7 days, with a range of 5 to 7 days, prior to both study nights. Bedtimes were recorded using daily sleep diaries and wrist actigraphy monitoring (Actiwatch Type AWL, CamNtech, Cambridge, UK).

Participants were permitted to consume a restricted amount (1 standard portion) of alcohol and caffeine up to 4 days before the experiment. Complete abstinence from alcohol and caffeine was required from that point until the day of the experiment. Participants documented their consumption of these substances and any medication through self-reports. On the day of the experiment, participants were instructed to refrain from activities known to affect sleep, such as sauna use or engaging in extreme sports.

A randomized, double-blind, crossover study design was employed, with each participant undergoing two experimental nights: a stimulation night (STIM) and a SHAM night. The two nights were separated by a minimum of 5 days.

On the days of the experiment, participants arrived at the laboratory at 6:50 p.m. and completed a day-report questionnaire. Following the preparation of EEG, EMG (electromyogram), and ECG (electrocardiogram) recordings, a 3-minute resting-state EEG was recorded. Subsequently, participants performed four behavioral tasks (word-pairs, object-locations, alertness, finger-tapping task) between 8:30 and 9:45 p.m., in the specified order. Task motivation was assessed before each task. The object-location task was introduced starting from the 2nd participant, leading to 13 complete data sets. The finger-tapping task was only introduced starting from the 8th participant, which is why the data were not included in the analysis.

Sleep EEG recordings commenced at 10:30 p.m. with lights off. Auditory stimulation was initiated after 10 minutes of stable NREM (non-rapid eye movement) sleep was detected automatically, specifically targeting sleep stages N2 and N3. Following an 8-hour sleep opportunity, lights were turned on at 6:30 a.m., and participants were immediately requested to complete a sleep report. At 7:15 a.m., which was 45 minutes after lights were turned on to minimize sleep inertia, resting-state EEG and all behavioral tasks were repeated. The total duration of the task session was approximately 1 hour. If participants indicated habitual earlier bedtimes, the schedule was advanced by 30 minutes.

Three weeks after each experimental night, the retrieval of both memory tests (word-pairs and object-locations) was repeated through video calls conducted with participants from their respective homes. This follow-up assessment allowed for the examination of long-term memory performance and retention beyond the immediate post-experimental period (data not included).

### Behavioral tasks

Four behavioral tasks were administered in the study. These tasks included two declarative memory tests, one focused on verbal memory, the other on spatial memory. Additionally, an alertness and a motor-learning task (data not included) were used to assess different cognitive functions and abilities.

#### Associated word-pair memory task

Auditory presentation of words was conducted using PsychoPy [version 2020.1.3; 60] through the loudspeakers of a laptop. To ensure test standardization, word-pairs were previously recorded by the same individual.

In the evening session, participants were tasked with learning a list of 40 word-pairs that were characterized by low emotionality and high tangibility (*encoding phase*, **Supplementary Fig. 5D**). Immediately following the encoding phase, participants were presented with only the first word of each word-pair and were required to recall the associated word within the pair without a specific time limit (*immediate retrieval*). If an incorrect response was provided, the full word-pair was presented again. The immediate retrieval process continued until participants successfully recalled at least 60% (24 word-pairs) of the word-pairs correctly. The experimenter determined whether participant’s spoken responses were correct or false through a button press, ensuring accurate calculation of the 60% threshold. A response was deemed correct regardless of its singular or plural form. However, synonyms were not accepted.

In the morning session, participants were presented with the first word of each word-pair and were asked to recall the associ-ated word (*delayed retrieval*). No feedback was provided during the delayed retrieval phase.

Afterwards, in the morning session, participants underwent a new encoding phase where they learned another set of 40 word-pairs (*new encoding*). Immediate retrieval of the newly learned word-pairs was conducted once without replaying falsely recalled word-pairs. No feedback was given during this process.

To assess long-term memory consolidation, delayed retrieval of word-pairs from the initial encoding phase in the evening session occurred three weeks after the experimental night. Participants performed the retrieval task from their homes via video call, and no feedback was provided during the retrieval session. This data is not presented.

Among the four lists (**Supplementary Fig. 6)**, lists 1 and 2 were consistently used during the evening encoding sessions, while lists 3 and 4 were exclusively used during the morning sessions. Specifically, list 1 was always followed by list 3, and list 2 was always followed by list 4. The word-pairs within each list were presented in a randomized order, while the assignment of lists was randomized across different conditions.

The inter-stimulus interval (ISI) between the words within a word-pair, starting from the voice play onset of the first word to the the onset of the second word, was set to 2 seconds. Additionally, the ISI between consecutive word-pairs, starting from the voice play onset of the second word to the onset of the first word of the next pair, was set to 5 seconds.

A pool of 160 word-pairs was utilized (**Supplementary Fig. 6)**, consisting of 112 word-pairs adopted from previous studies (32 from Marshall et al. 61, 80 from Wilhelm et al. 62, and 48 newly selected word-pairs from Hager and Hasselhorn 63).

#### Object-location task

The object-pairs were assigned specific locations on a 5 × 6 checkerboard-pattern, consisting of 30 gray tiles separated by a white grid (**Supplementary Fig. 5E**). The experiment was implemented using Psychopy [version 2020.1.3; 60].

During the evening session, participants were instructed to learn the locations of 15 neutral day-to-day object-pairs (encoding phase). During the encoding phase, the first object of a pair was presented for a duration of 1 second. Subsequently, the second object of the pair was revealed, allowing participants to view the entire object-pair for an additional 3 seconds. The ISI between each object-pair presentation was set to 3 seconds.

The encoding process was conducted twice, with each presentation using a different randomized order of object-pairs. A separate practice session was not required. After encoding, participants have seen the location of each object-pair twice.

Following the encoding phase, participants were presented with only the first object and were required to click on the gray square corresponding to the location of the corresponding object-pair (immediate retrieval). There was no specific time limit imposed during this task. The location of the correct object-pair was revealed regardless of the response accuracy. Immediate retrieval continued until participants successfully recalled at least 60% (9 object-pairs) correctly.

In the morning session, participants were presented with the first object and were asked to click on the gray square representing the corresponding location of the object-pair (delayed retrieval). The location of the correct object-pair was not revealed to prevent additional encoding. Instead, the selected tile was highlighted by changing the color of the tile edges from white to cyan. Delayed retrieval was repeated 3 weeks after the experimental night from home via video-call. Tiles were numbered so that participants verbally expressed their response to the experimenter who then clicked on the corresponding tile. This data is not presented.

All 30 object-pairs utilized in this study (**Supplementary Fig. 7)** were adopted from a previous study [64].

#### Alterness task

Both in the evening and morning sessions, participants completed the alertness subtest of the Test of Attentional Performance (TAP) software [version 2.3.1; 65]. The alertness task assessed reaction times in two different conditions to evaluate both tonic and phasic alertness.

Tonic alertness was evaluated by measuring the reaction time (button press) in response to the appearance of a white cross on a black screen. Phasic alertness, on the other hand, was assessed by announcing the impending appearance of the white cross with a beep sound that was presented shortly before (with a random lag time).

The alertness task consisted of four blocks, with two blocks dedicated to assessing tonic alertness and the other two blocks focused on phasic alertness. Each block comprised 20 trials (tonic - phasic - phasic - tonic), resulting in a total of 80 trials across the entire task.

#### Finger sequence tapping task

Although the specific data from this test is not presented here due to the low number of participants, we will provide an explanation of the procedure for the sake of protocol completion.

In the evening session, participants engaged in a computerized finger sequence tapping task [adapted from 27], and implemented using Psychopy [version 2020.1.3; 60]. During the training trial, participants were instructed to type a simple sequence repeatedly for 30 seconds, aiming to complete as many repetitions as possible. Following the training trial, a 30-second rest period was given to prevent fatigue. Subsequently, participants completed 12 consecutive trials of typing the evening sequence. Each trial lasted 30 seconds and was followed by a 30-second rest period.

In the morning session, participants performed a new sequence for 12 consecutive trials, each lasting 30 seconds and separated by a 30-second rest period. Following the morning sequence, participants returned to type the evening sequence for 3 consecutive trials, with each trial lasting 30 seconds and separated by a 30-second rest period.

During the finger sequence tapping task, participants viewed the sequence on the screen, where the numbers 1, 2, 3, and 4 represented the index, middle, ring, and little finger of the right hand, respectively. As participants executed the sequence by tapping the corresponding keys, a black dot would appear below the current number on the screen to indicate that a response was recorded. However, no accuracy feedback was given during the task.

Upon completing a sequence, the screen was refreshed, displaying the same sequence, but with no black dots present. The training sequence consisted of the pattern 1-1-2-2-3-4. For the evening sequences, participants performed two different patterns: 2-4-1-3-1-2 and 2-1-3-1-4-2. In the morning session, two additional sequences were performed: 4-2-3-1-4-3 and 3-4-1-3-2-4.

### Sleep hd-EEG recordings

During both experimental nights, hd-EEG data was recorded using an Electrical Geodesics Sensor Net for long-term monitoring (128 channels, Net Amps 400 series, Electrical Geodesics Inc., EGI, Eugene, OR) and acquired using EGI Net Station version 5.4. Electrooculographic (EOG) and submental electromyographic (EMG) signals were utilized for manual sleep scoring. To ensure optimal recording conditions, two additional electrodes (gold, Grass Technologies, West Warwick, RI) were attached to the earlobes and served as alternative references if necessary.

Electrocardiogram (ECG) electrodes (oval solid gel, Skintact, Innsbruck, AT) were positioned two fingers below the right clavicle and two fingers below the lowest left rib. After proper adjustment of the net to the vertex and mastoids, the skin underneath each electrode was prepared, and an electrolyte gel (ECI Electro-Gel, Electro-Cap International, Inc., Eaton, OH) was applied to ensure high conductance and maintain signal quality throughout the night. The impedance of the EEG electrodes was initially kept below 50 kΩ (5 kΩ for gold electrodes) at the start of the recording and rechecked in the morning. During recording, all channels were referenced to Cz and sampled at a rate of 500 Hz. The ECG was recorded using a bipolar reference configuration.

Subsequently, the EEG data was exported using EGI Net Station version 5.4, applying a specific 0.1 Hz high-pass filter to remove slow drifts and voltage jumps while ensuring the preservation of relevant frequency information.

### Auditory stimulation

For real-time slow-wave detection and phase-locked auditory stimulation, a configurable mobile EEG system [referenced as MHSL-SleepBand in 66] was utilized. Stimulation triggers from the MHSL-SleepBand were sent as digital input to the EGI amplifier, allowing their integration into the hd-EEG signal.

Regarding the placement of electrodes for the MHSL-SleepBand, a gold electrode from Grass Technologies (West Warwick, RI) was positioned adjacent to C3, located between electrodes 29, 30, and 36 of the hd-EEG net, enabling real-time monitoring of EEG activity. Two additional gold electrodes were attached to the mastoids, serving as the ground (ipsilateral to the detection electrode) and the reference (contralateral to the detection electrode).

For the delivery of auditory stimuli, participants had on-ear headphones (sleepPhones, AcousticSheep LLC, Peninsula Drive, Pennsylvania) securely taped to their ears.

Auditory stimuli were presented at 50 dB. The loudness was verified using a sound level meter (UNI-T UT352 Type 2, Uni-Trend Technology EU GmbH, Augsburg, Germany). The measurement was taken with the sound level meter placed directly against the headphones. Based on subjective perception, stimuli delivered via headphones were clearly audible.

A comprehensive description of the MHSL-SleepBand algorithm has been published previously [66]. In summary, the algorithm utilized three parallel binary classifiers:

1. *Sleep detection classifier:* This classifier continuously assessed increased SWA, reduced beta EEG activity, as well as the ratio of the two from the past 80 s of EEG to detect specific sleep stages, namely N2+N3 sleep. Sleep detection was satisfied when the predefined power thresholds were met.
2. *SWA classifier:* The SWA classifier continuously evaluated the SWA in the past 4 s of EEG signal. It aimed to detect as many slow waves as possible. When a predefined power threshold was met, SWA detection was satisfied.
3. *Phase detection classifier:* This classifier employed a phase-locked loop (PLL) architecture to estimate the phase of the input EEG signal. Its purpose was to identify the occurrence of 1 Hz waves and present auditory stimuli shortly before the negative peak of these slow waves.

When all three conditions were satisfied, an auditory stimulus in the form of a 50 ms tone of 1/f pink noise was delivered through headphones. The applied thresholds were optimized for the specific age group and electrode location by conducting simulations to evaluate algorithm performance based on previously collected data.

During the stimulation nights, auditory stimuli were presented during 16 second ON windows. These windows provided the opportunity for the delivery of auditory stimulation. In contrast, 8 second OFF windows were implemented to withhold auditory stimulation. ON and OFF windows for stimulus presentation were initiated when the sleep detection classifier criteria were met and discontinued whenever the criteria were no longer fulfilled, as a measure for battery conservation. Stimulation was applied throughout the entire night, starting after the detection of 10 minutes of stable NREM sleep. This ensured that auditory stimuli were administered during the targeted sleep stages of interest. In the SHAM night, the same timing flags for stimulation were recorded alongside the EEG data, but no actual auditory stimuli were presented.

### EEG preprocessing & sleep scoring

All EEG processing steps and analyses were performed in Matlab (R2017b, R2021b, R2022a, R2022b & R2023a; The Math-Works, Inc., Natick, Massachusetts) and EEGLAB [version 2021.1; 67]. Sleep scoring was performed using a software obtained from the Institute of Pharmacology and Toxicology of the University of Zurich, Switzerland.

#### Filters & down-sampling

The EEG data underwent several processing steps (**Supplementary Fig. 4)**. First, the EEG data was subjected to a low-pass filter with a cut-off of −6 dB at 39.86 Hz (filter order: 92 at a sampling rate of 500 Hz). Subsequently, the data was down-sampled to a sampling rate of 125 Hz and high-pass filtered with a cut-off of −6 dB at 0.37 Hz (filter order: 1494 at 125 Hz).

In recordings with strong sweat artifacts, as well as all corresponding recorded nights of the same participant, the EEG data was subjected to a stronger high-pass filter to ensure accurate sleep scoring. The high-pass filter had a cut-off of −6 dB at 0.75 Hz (filter order: 1246 at 125 Hz). This occurred in four out of 14 participants. The filter specifically attenuates frequencies below 0.9 Hz, with increasing attenuation down to 0.6 Hz, below which frequencies are nearly completely attenuated. Therefore, all analyses related to frequency in our study are focused on frequencies above 1 Hz.

All filters, including those mentioned earlier and any subsequent filters, were Kaiser-window based FIR filters and were applied in a one-way manner with zero-phase shift, ensuring that filtering did not introduce any phase distortion. The filter order (in samples) is directly related to the sampling rate of the EEG, meaning that doubling the sampling rate would require doubling the filter order as well.

While zero-phase filtering corrects the linear phase delay induced by the applied filter, thus facilitating the interpretation of the relationship between ERP waveforms and cross-frequency effects in ERSP analyses, the non-causal nature of the filter may inadvertently project post-stimulus effects into pre-stimulus periods. Hence, caution is advised when interpreting potential prestimulus differences between conditions.

#### Sleep scoring

Sleep stages were manually scored in 20-second epochs by one sleep expert and verified by another sleep expert using EEG, EOG, and EMG signals. The scoring was conducted according to standard criteria [68], considering five vigilance stages: wake, N1, N2, N3, and REM sleep. Both scorers were blinded to the experimental condition and unaware of which night belonged to each participant. However, they were aware which nights belonged to the same participant.

For sleep scoring, the EEG data was referenced to the contralateral mastoid and down-sampled to 128 instead of 125 Hz to match the requirements of the scoring software.

The EOG data was high-pass filtered with a cut-off of −6 dB at 0.13 Hz (filter order: 1594 at 128 Hz) to preserve slow-rolling eye movements. The EMG data was low-pass filtered with a cut-off of −6 dB at 96.01 Hz (filter order: 230 at 500 Hz), notch filtered at 47.49 Hz (filter order: 368 at 500 Hz), down-sampled to 128 Hz, and high-pass filtered with a cut-off of −6 dB at 14.01 Hz (filter order: 64 at 128 Hz).

#### Artifact removal

During the manual sleep scoring process, any epoch containing artifacts in the channels visible during sleep scoring were excluded from further analyses. Afterward, 20-second epochs from 126 EEG channels (excluding the 2 submental EMG channels) were semi-automatically assessed using a custom graphical user interface (GUI) designed specifically for sleep hd-EEG data [69]. In short, the raw EEG signal, as well as the distribution and topography of four sleep quality markers, were examined to detect artifactual NREM (N1+N2+N3) epochs. If more than two neighboring channels exhibited artifacts, the respective epoch was considered artifactual and removed. All remaining epochs were retained for subsequent EEG preprocessing steps.

#### Epoch-wise interpolation

Any channels marked with artifacts were subjected to interpolation in the affected epochs specifically using the *interpEPO()* function [69]. As stated previously, only epochs in which no more than two neighboring channels contained artifacts were considered for interpolation. Following this procedure, of all the scored NREM (N2+N3) epochs, on average, 35.00% (SD = 30.96%, ranging from 2.50% to 96.34%) could be interpolated, while 6.77% (SD = 3.26%, ranging from 2.79% to 17.50%) were removed. The remaining epochs were free of artifacts.

#### Re-referencing

Unless specified otherwise, the EEG data was referenced to the mean of all 110 channels located above the ear (**Supplementary Fig. 8)**. Average referencing offers the advantage of not favoring any particular cortical location, thereby facilitating the identification of local effects [e.g., 70]. When local effects were not of primary interest, linked mastoid reference was used.

### Analysis of sleep EEG

All subsequent analyses were performed exclusively on artifact-free NREM (N2+N3) epochs or ON|OFF windows. Functions for circular outcome measures (e.g., mean phase) were used from the toolbox for circular statistics [71]. The mean of all channels and the channel average refer to the mean of the 110 channel above the ear if not stated otherwise.

#### Auditory evoked response (AER)

The EEG data, referenced to the linked mastoids, was aligned to isolated stimuli and a short time-window (−0.4 – 3 s) was extracted. The EEG data within the selected time-windows was averaged across trials (stimuli) for a given channel and sample point, resulting in averaged waveforms after both STIM and SHAM flags. The AER is the difference in waveforms between conditions, thereby subtracting the underlying phase-locked slow wave (mean aveform in SHAM) from the AER that is convoluted with the phase-locked slow wave (mean waveform in STIM). The AER was computed separately for each participant and then averaged across participants.

#### Topography of N550

For each night, the latency of the N550 component was determined individually. This was done by identifying the latency of minimum value in the ERP waveform within a specific time-window (0.35 – 0.7 s) following the presentation of the isolated stimuli. The ERP consisted of the average signal from channels within a predefined frontal ROI.

Once the latency of the minimum value was identified, the corresponding amplitude at that latency was extracted in all channels. This amplitude represented the amplitude of the N550 component in a given channel.

#### Time-frequency analyses

Time-frequency analyses were performed using Morlet wavelets with 3 cycles at the lowest (1 Hz) to 10 cycles at the highest frequency (25 Hz; logarithmically spaced), resulting in phase and spectral power (event-related spectral perturbation; ERSP) values for 49 frequencies between 1 and 25 Hz (in steps of 0.5 Hz). The average referenced EEG data was aligned to the onset of isolated stimuli. For a given ON window, EEG data within a short time-window (−3.2 – 5.608 s) was used as input for a standard wavelet transformation routine [72]. The first and last 3 s of data were treated as a buffer zone to prevent edge artifacts, resulting in phase and spectral power values for 351 time points between −0.2 and 2.6 s (at 125 Hz). To minimize file sizes, phase and spectral power values were subsequently down-sampled to 50 Hz. To eliminate participant and frequency biases, spectral power values were normalized by dividing them by the average spectral power of the entire time window from the average of both nights (participant- and frequency-wise).

#### Spindle detection

Spindles were detected using a previously described method [42]. In short, the EEG data was band-pass filtered using a Chebyshev Type II filter with cut-offs of −6 dB at 12.03 and 17.05 Hz (filter order: 2). Then, within artifact-free non-REM sleep epochs, spindles were identified whenever the amplitude fluctuation exceeded five times the average amplitude of a specific channel. Of all the identified spindles, those were selected which had a freuency between 11 and 16 Hz, had a duration between 0.5 and 2 s, and did not exceed a maximum amplitude of 100 *µ*V. Whenever the point of absolute maximum amplitude of the spindle occurred within two seconds following isolated stimuli, the phase of the underlying slow wave (at frequencies between 1 and 4 Hz, in steps of 1 Hz) was extracted at this point. Finally, for a given channel, the number of spindles occurring within the positive (up-phase coupled) or negative half-wave (down-phase coupled) of a slow wave was counted.

#### Power spectral density (PSD)

For each night, PSD values were calculated from average referenced EEG data. The PSD was computed using Matlab’s *pwelch()* function (Welch method with 4-second Hanning windows, 50% overlap, and a frequency resolution of 0.25 Hz).

To compute overnight PSD values, EEG data within artifact-free N2, N3, or NREM (N2+N3) epochs were utilized. This resulted in separate overnight PSD values for N2 epochs, N3 epochs, and combined NREM epochs (N2+N3). Each epoch was divided into nine overlapping 4-second windows.

For PSD within ON|OFF windows, ON windows were divided into smaller segments (ON1, ON2, OFF), each with a duration of 8 seconds. Each segment was divided into three overlapping 4-second windows. Only ON|OFF windows that were entirely located within artifact-free NREM epochs were included in the analysis.

#### Overnight SWA

First, PSD values were averaged over epochs, separately for each frequency and channel (**Supplementary Fig. 3C**). For the computation of SWA, the lower boundary for SWA was set at 1 Hz due to the presence of sweat artifacts in some recordings. To obtain overnight SWA, averaged PSD values within the frequencies of interest (1 – 4.5 Hz) were summed and multiplied by the frequency resolution (0.25 Hz), separately for each channel (**Supplementary Fig. 3A**).

#### SWA within ON|OFF windows

For SWA within ON|OFF windows, PSD values were averaged over specific 8-second segments that corresponded to each ON|OFF window. ON|OFF windows were divided into three subsets based on the number of stimuli within the first 6 seconds of ON windows. These subsets corresponded to windows with few (1 – 2), medium (3 – 4), or many (5 – 8) stimuli, resulting in nine different window types (ON1, ON2, and OFF windows with either few, medium, or many stimuli).

For each window type, PSD values were averaged over windows, separately for each frequency and channel. SWA was calculated as described for overnight SWA. Percentage change between conditions was computed by taking the difference in SWA (STIM−SHAM) and dividing it by the average SWA levels of both nights (channel-wise).

#### Relative PSD within ON|OFF windows

Relative PSD values describe a redistribution (a shift) of PSD values within ON|OFF windows. To obtain relative PSD values, PSD values were divided by the channel average PSD of the entire ON|OFF window, separately for each frequency, window, and channel. For each of the nine window types, the mean of all channels and windows was computed to obtain average relative PSD values.

#### Phase estimation of auditory stimuli

To estimate the phase precision of presented stimuli (**Supplementary Fig. 2C**), the EEG data underwent filtering within the slow-wave range. For that, the EEG data was subjected to a low-pass filter with a cut-off of −6 dB at 5.00 Hz (filter order = 904 at 500 Hz), down-sampled to 125 Hz, and high-pass filtered with a cut-off of −6 dB at 0.37 Hz (filter order = 1494 at 125 Hz). The EEG data was referenced to the linked mastoids and the first and last 14.940 samples (10 times the filter order) were set to zero. This step was performed to mitigate edge artifacts that could distort the Hilbert transform. Phase values were extracted by applying the *hilbert()* function to the EEG data of each channel. Phase values corresponding to the times of stimulation within artifact-free NREM (N2+N3) epochs were extracted and stored for further analysis.

#### Phase bins

For each participant, phase values were clustered into one of ten 30° bins (**Supplementary Fig. 2A**). This binning process involved using the mean phase value of channels 29, 30, and 36 (which are around the target electrode) with the help of the *circ_mean()* function. The percentage of values falling within each specific phase bin was then calculated.

#### Phase variance

The circular standard deviation of phase values was computed using the *circ_std()* function of the circular statistics toolbox, separately for each channel (**Supplementary Fig. 2B**).

### Analysis of behavioral tasks

Due to non-compliance with task instructions, one participant was excluded from the behavioral analyses. As a result, there were 13 complete datasets available for the analysis of the word-pair and alertness task, and 12 complete datasets for the object-location task.

#### Analysis of memory tasks

The number of correctly recalled word-pairs (object-pairs) in the evening and morning were used to assess overnight consolidation. The measure of overnight consolidation was computed as the difference in the number of correctly recalled items from morning to evening. Additionally, the number of correctly recalled word-pairs from the new list served as a measure of encoding capabilities the next morning. This is because the immediate retrieval of a newly learned word list on the subsequent morning is solely influenced by encoding and retrieval capabilities rather than overnight consolidation.

In case participants required several trials to accomplish the 60% criterion, the number of correctly recalled items comprised the total number of correctly recalled items across trials.

#### Analysis of alertness task

Reaction times (RT, in ms) were averaged across trials within each block. For both phasic and tonic alertness, the average RTs were computed across blocks, along with their standard deviation. These measures provide an indication of alertness and sensitivity to sleep pressure [73, 74]. Overnight performance change was assessed by calculating the difference in average RT and the standard deviation of RT from morning to evening.

#### Correlations

Spearman’s rank correlation was used to examine the relationship between the performance in overnight memory consolidation of the associated word-pair task and the quantity of detected spindles following isolated stimuli (STIM) or their respective stimulation flags (SHAM). Specifically, channel-wise correlations were performed on the number of stimulus-associated spindles, both as absolute values and normalized to the total number of overnight spindles. Furthermore, correlations were performed on the normalized number of spindles coupled to the up-phase or down-phase of slow waves, as well as on the difference in the normalized number of up-phase and down-phase coupled spindles.

### Statistics

Statistics for EEG data and correlations were performed in MATLAB (R2022a, R2022b, R2023a, The Math-Works, Inc., Natick, Massachusetts). Statistics for behavioral data were performed in R (4.2.1) and RStudio (2022.07.1).

#### Topographical comparisons

For topographical comparisons between conditions (STIM−SHAM) of SWA, the number of detected spindles, and the amplitude of the N550 component, paired Student’s t-tests (*α* = .05, two-tailed) were performed (channel-wise). For topographical correlations between two variables, Spearman’s rank correlation coefficients were computed (*α* = .05, two-tailed). To account for multiple comparisons, non-parametric cluster-based statistical mapping was applied [75].

#### Time-frequency comparisons

ERSP comparisons between conditions (STIM−SHAM) were examined using paired Student’s t-tests (*α* = .0001, two-tailed; pixel-wise). To account for multiple comparisons, non-parametric cluster-based statistical mapping was applied.

#### Non-parametric cluster-based statistical mapping

In short, the condition label (STIM or SHAM) was pseudorandomly permuted across participants, so that participants pseudo-randomly switched conditions. The data was permuted 5000 times. The number of permutations is restricted by the number of participants (2^*N*^ − 1). Each permutation resulted in mutually exclusive condition labels. Paired Student’s t-tests were performed in each permutation and the maximum cluster size of significant neighboring electrodes (or pixels) was computed, separately for positive and negative t-values. This resulted in two distributions of cluster sizes, one for positive and one for negative t-values, with as many values as permutations performed.

For topographical comparisons, the 97.5th percentile in each distribution, for the ERSP, the 99.99th percentile in each distribution was defined as the critical cluster size threshold, respectively, accounting for the higher number of comparisons in ERSP analyses due to the higher number of pixels involved. When the original data showed a cluster of significant electrodes (or pixels) equal or larger than the critical cluster size threshold, this cluster of electrodes (or pixels) was considered significant. Multiple clusters could co-exist.

#### AER comparisons

AER comparisons between conditions (STIM−SHAM) were performed at each sample point using paired Student’s t-tests (*α* = .05, two-tailed; sample-wise). False discovery rate correction [76] was applied to account for multiple comparisons.

#### Behavioral comparisons

Comparisons of all behavioral measures between conditions (STIM−SHAM) were performed using paired Student’s t-tests (*α* = .05, two-tailed).

## References

[1] Chiara Cirelli and Giulio Tononi. The why and how of sleep-dependent synaptic down-selection. In Seminars in cell & developmental biology, volume 125, pages 91–100. Elsevier, 2022.

[2] Natalie L Hauglund, Chiara Pavan, and Maiken Nedergaard. Cleaning the sleeping brain–the potential restorative function of the glymphatic system. Current Opinion in Physiology, 15:1–6, 2020.

[3] Susanne Diekelmann and Jan Born. The memory function of sleep. Nature Reviews Neuroscience, 11(2):114–126, 2010.

[4] Johannes Vosskuhl, Daniel Strüber, and Christoph S Herrmann. Non-invasive brain stimulation: a paradigm shift in understanding brain oscillations. Frontiers in human neuroscience, 12:211, 2018.

[5] Jens G Klinzing, Niels Niethard, and Jan Born. Mechanisms of systems memory consolidation during sleep. Nature neuroscience, 22(10):1598–1610, 2019.

[6] Daisuke Miyamoto, Daichi Hirai, and Masanori Murayama. The roles of cortical slow waves in synaptic plasticity and memory consolidation. Frontiers in neural circuits, 11:92, 2017.

[7] Giulio Tononi and Chiara Cirelli. Sleep and the price of plasticity: from synaptic and cellular homeostasis to memory consolidation and integration. Neuron, 81(1): 12–34, 2014.

[8] Yo-El S Ju, Sharon J Ooms, Courtney Sutphen, Shannon L Macauley, Margaret A Zangrilli, Gina Jerome, Anne M Fagan, Emmanuel Mignot, John M Zempel, Jurgen AHR Claassen, et al. Slow wave sleep disruption increases cerebrospinal fluid amyloid-β levels. Brain, 140(8):2104–2111, 2017.

[9] Hong-Viet V Ngo, Thomas Martinetz, Jan Born, and Matthias Mölle. Auditory closed-loop stimulation of the sleep slow oscillation enhances memory. Neuron, 78(3):545–553, 2013.

[10] Hong-Viet V Ngo, Arjan Miedema, Isabel Faude, Thomas Martinetz, Matthias Mölle, and Jan Born. Driving sleep slow oscillations by auditory closed-loop stimulation—a self-limiting process. Journal of Neuroscience, 35(17):6630–6638, 2015.

[11] Ju Lynn Ong, June C Lo, Nicholas IYN Chee, Giovanni Santostasi, Ken A Paller, Phyllis C Zee, and Michael WL Chee. Effects of phase-locked acoustic stimulation during a nap on eeg spectra and declarative memory consolidation. Sleep medicine, 20:88–97, 2016.

[12] Miika M Leminen, Jussi Virkkala, Emma Saure, Teemu Paajanen, Phyllis C Zee, Giovanni Santostasi, Christer Hublin, Kiti Müller, Tarja Porkka-Heiskanen, Minna Huotilainen, et al. Enhanced memory consolidation via automatic sound stimulation during non-rem sleep. Sleep, 40(3), 2017.

[13] Nelly A Papalambros, Giovanni Santostasi, Roneil G Malkani, Rosemary Braun, Sandra Weintraub, Ken A Paller, and Phyllis C Zee. Acoustic enhancement of sleep slow oscillations and concomitant memory improvement in older adults. Frontiers in human neuroscience, 11:109, 2017.

[14] Ju Lynn Ong, Amiya Patanaik, Nicholas IYN Chee, Xuan Kai Lee, Jia-Hou Poh, and Michael WL Chee. Auditory stimulation of sleep slow oscillations modulates subsequent memory encoding through altered hippocampal function. Sleep, 41(5): zsy031, 2018.

[15] Simon Henin, Helen Borges, Anita Shankar, Cansu Sarac, Lucia Melloni, Daniel Friedman, Adeen Flinker, Lucas C Parra, Gyorgy Buzsaki, Orrin Devinsky, et al. Closed-loop acoustic stimulation enhances sleep oscillations but not memory performance. eneuro, 6(6), 2019.

[16] Elena Krugliakova, Jelena Skorucak, Georgia Sousouri, Sven Leach, Sophia Snipes, Maria Laura Ferster, Giulia Da Poian, Walter Karlen, and Reto Huber. Boosting recovery during sleep by means of auditory stimulation. Frontiers in neuroscience, 16, 2022.

[17] Elena Krugliakova, Carina Volk, Valeria Jaramillo, Georgia Sousouri, and Reto Huber. Changes in cross-frequency coupling following closed-loop auditory stimulation in non-rapid eye movement sleep. 10(1):1–12, 2020.

[18] Nelly A Papalambros, Sandra Weintraub, Tammy Chen, Daniela Grimaldi, Giovanni Santostasi, Ken A Paller, Phyllis C Zee, and Roneil G Malkani. Acoustic enhancement of sleep slow oscillations in mild cognitive impairment. Annals of clinical and translational neurology, 6(7):1191–1201, 2019.

[19] Charmaine Diep, Suzanne Ftouni, Jessica E Manousakis, Christian L Nicholas, Sean PA Drummond, and Clare Anderson. Acoustic slow wave sleep enhancement via a novel, automated device improves executive function in middle-aged men. Sleep, 43(1):zsz197, 2020.

[20] Jules Schneider, Penelope A Lewis, Dominik Koester, Jan Born, and Hong-Viet V Ngo. Susceptibility to auditory closed-loop stimulation of sleep slow oscillations changes with age. Sleep, 43(12):zsaa111, 2020.

[21] Alexander Prehn-Kristensen, Hong-Viet V Ngo, Luisa Lentfer, Julia Berghäuser, Lena Brandes, Larissa Schulze, Robert Göder, Matthias Mölle, and Lioba Baving. Acoustic closed-loop stimulation during sleep improves consolidation of reward-related memory information in healthy children but not in children with attention-deficit hyperactivity disorder. Sleep, 43(8):zsaa017, 2020.

[22] Marcus O Harrington, Hong-Viet V Ngo, and Scott A Cairney. No benefit of auditory closed-loop stimulation on memory for semantically-incongruent associations. Neurobiology of learning and memory, 183:107482, 2021.

[23] Ping Koo-Poeggel, Soé Neuwerk Eike Petersen, Jan Grasshoff, Matthias Mölle, Thomas Martinetz, and Lisa Marshall. Closed-loop acoustic stimulation during an afternoon nap to modulate subsequent encoding. Journal of Sleep Research, 31 (6):e13734, 2022.

[24] Stephanie Huwiler, Manuel Carro Dominguez, Silja Huwyler, Luca Kiener, Fabia Stich, Rossella Sala, Florent Aziri, Anna Trippel, Christian Schmied, Reto Huber, et al. Effects of auditory sleep modulation approaches on brain oscillatory and cardiovascular dynamics. bioRxiv, 2022.

[25] Caroline Lustenberger, M Laura Ferster, Stephanie Huwiler, Luzius Brogli, Esther Werth, Reto Huber, and Walter Karlen. Auditory deep sleep stimulation in older adults at home: a randomized crossover trial. Communications Medicine, 2(1): 1–16, 2022.

[26] Carlos G Moreira, Christian R Baumann, Maurizio Scandella, Sergio I Nemirovsky, Sven Leach, Reto Huber, and Daniela Noain. Closed-loop auditory stimulation method to modulate sleep slow waves and motor learning performance in rats. Elife, 10:e68043, 2021.

[27] Sara Fattinger, Toon T de Beukelaar, Kathy L Ruddy, Carina Volk, Natalie C Heyse, Joshua A Herbst, Richard HR Hahnloser, Nicole Wenderoth, and Reto Huber. Deep sleep maintains learning efficiency of the human brain. Nature communications, 8 (1):1–14, 2017.

[28] Marcus O Harrington and Scott A Cairney. Sounding it out: Auditory stimulation and overnight memory processing. Current Sleep Medicine Reports, 7(3):112–119, 2021.

[29] Yujie Zhang and Reut Gruber. Focus: attention science: can slow-wave sleep enhancement improve memory? a review of current approaches and cognitive outcomes. The Yale journal of biology and medicine, 92(1):63, 2019.

[30] Vincenzo Crunelli, Magor L Lőrincz, Adam C Errington, and Stuart W Hughes. Activity of cortical and thalamic neurons during the slow (< 1 hz) rhythm in the mouse in vivo. Pflügers Archiv-European Journal of Physiology, 463(1):73–88, 2012.

[31] Georgia Sousouri, Elena Krugliakova, Jelena Skorucak, Sven Leach, Sophia Snipes, Maria Laura Ferster, Giulia Da Poian, Walter Karlen, and Reto Huber. Neuromodulation by means of phase-locked auditory stimulation affects key marker of excitability and connectivity during sleep. Sleep, 45(1):zsab204, 2022.

[32] P Halasz. Arousals without awakening—dynamic aspect of sleep. Physiology & behavior, 54(4):795–802, 1993.

[33] Célyne H Bastien, Cécile Ladouceur, and Kenneth B Campbell. Eeg characteristics prior to and following the evoked k-complex. Canadian Journal of Experimental Psychology/Revue canadienne de psychologie expérimentale, 54(4):255, 2000.

[34] Martin Roth, John Shaw, and Joy Green. The form, voltage distribution and physiological significance of the k-complex. Electroencephalography and clinical neurophysiology, 8(3):385–402, 1956.

[35] John Gora, Ian M Colrain, and John Trinder. The investigation of k-complex and vertex sharp wave activity in response to mid-inspiratory occlusions and complete obstructions to breathing during nrem sleep. Sleep, 24(1):81–89, 2001.

[36] Kimberly A Cote, Duncan R De Lugt, Susan D Langley, and Kenneth B Campbell. Scalp topography of the auditory evoked k-complex in stage 2 and slow wave sleep. Journal of Sleep Research, 8(4):263–272, 1999.

[37] Célyne H Bastien, Kate E Crowley, and Ian M Colrain. Evoked potential components unique to non-rem sleep: relationship to evoked k-complexes and vertex sharp waves. International Journal of Psychophysiology, 46(3):257–274, 2002.

[38] Florin Amzica and Mircea Steriade. The k-complex: its slow (< 1-hz) rhythmicity and relation to delta waves. Neurology, 49(4):952–959, 1997.

[39] Vasileios Kokkinos, Andreas M Koupparis, and George K Kostopoulos. An intra-k-complex oscillation with independent and labile frequency and topography in nrem sleep. Frontiers in human neuroscience, 7:163, 2013.

[40] B Kurella, A Heitmann, M Golz, and U Dormann. The probability of eliciting k-complexes during sleep. J Sleep Res, 1(suppl 1):124, 1992.

[41] R Landwehr. Detection of activation phases and quantification of coupling in nrem sleep eeg by pointwise transinformation. Sleep Medicine, 8(1):65–72, 2007.

[42] Fabio Ferrarelli, Reto Huber, Michael J Peterson, Marcello Massimini, Michael Murphy, Brady A Riedner, Adam Watson, Pietro Bria, and Giulio Tononi. Reduced sleep spindle activity in schizophrenia patients. American Journal of Psychiatry, 164(3):483–492, 2007.

[43] Dag Neckelmann and Reidun Ursin. Sleep stages and eeg power spectrum in relation to acoustical stimulus arousal threshold in the rat. Sleep, 16(5):467–477, 1993.

[44] Randolph F Helfrich, Janna D Lendner, and Robert T Knight. Aperiodic sleep networks promote memory consolidation. Trends in Cognitive Sciences, 25(8):648– 659, 2021.

[45] Jens G Klinzing, Matthias Mölle, Frederik Weber, Gernot Supp, Jörg F Hipp, Andreas K Engel, and Jan Born. Spindle activity phase-locked to sleep slow oscillations. Neuroimage, 134:607–616, 2016.

[46] Caroline Lustenberger, Flavia Wehrle, Laura Tüshaus, Peter Achermann, and Reto Huber. The multidimensional aspects of sleep spindles and their relationship to word-pair memory consolidation. Sleep, 38(7):1093–1103, 2015.

[47] Hermann Griessenberger, Dominik PJ Heib, Julia Lechinger, Nikolina Luketina, Marit Petzka, Tina Moeckel, Kerstin Hoedlmoser, and Manuel Schabus. Susceptibility to declarative memory interference is pronounced in primary insomnia. PloS one, 8(2):e57394, 2013.

[48] Manuel Schabus, Kerstin Hoedlmoser, Thomas Pecherstorfer, Peter Anderer, Georg Gruber, Silvia Parapatics, Cornelia Sauter, Gerhard Kloesch, Wolfgang Klimesch, Bernd Saletu, et al. Interindividual sleep spindle differences and their relation to learning-related enhancements. Brain research, 1191:127–135, 2008.

[49] Carmen E Westerberg, Bryce A Mander, Susan M Florczak, Sandra Weintraub, M-Marsel Mesulam, Phyllis C Zee, and Ken A Paller. Concurrent impairments in sleep and memory in amnestic mild cognitive impairment. Journal of the International Neuropsychological Society, 18(3):490–500, 2012.

[50] Matthew A Tucker and William Fishbein. The impact of sleep duration and subject intelligence on declarative and motor memory performance: how much is enough? Journal of sleep research, 18(3):304–312, 2009.

[51] Caroline Lustenberger, Angelina Maric, Roland Dürr, Peter Achermann, and Reto Huber. Triangular relationship between sleep spindle activity, general cognitive ability and the efficiency of declarative learning. PloS one, 7(11):e49561, 2012.

[52] Laura MJ Fernandez and Anita Lüthi. Sleep spindles: mechanisms and functions. Physiological reviews, 100(2):805–868, 2020.

[53] Matthew P Walker. A refined model of sleep and the time course of memory formation. Behavioral and brain sciences, 28(1):51–64, 2005.

[54] Anita Lüthi. Sleep spindles: where they come from, what they do. The Neuroscientist, 20(3):243–256, 2014.

[55] Ian M Colrain. The k-complex: a 7-decade history. Sleep, 28(2):255–273, 2005.

[56] Timothy J Teyler and Pascal DiScenna. The hippocampal memory indexing theory. Behavioral neuroscience, 100(2):147, 1986.

[57] Kaori Takehara-Nishiuchi. Neurobiology of systems memory consolidation. European Journal of Neuroscience, 54(8):6850–6863, 2021.

[58] Lisa Genzel, Marijn CW Kroes, Martin Dresler, and Francesco P Battaglia. Light sleep versus slow wave sleep in memory consolidation: a question of global versus local processes? Trends in neurosciences, 37(1):10–19, 2014.

[59] Péter Halász. The k-complex as a special reactive sleep slow wave–a theoretical update. Sleep Medicine Reviews, 29:34–40, 2016.

[60] Jonathan Peirce, Jeremy R Gray, Sol Simpson, Michael MacAskill, Richard Höchenberger, Hiroyuki Sogo, Erik Kastman, and Jonas Kristoffer Lindeløv. Psychopy2: Experiments in behavior made easy. Behavior research methods, 51(1):195–203, 2019.

[61] Lisa Marshall, Halla Helgadóttir, Matthias Mölle, and Jan Born. Boosting slow oscillations during sleep potentiates memory. Nature, 444(7119):610–613, 2006.

[62] Ines Wilhelm, Susanne Diekelmann, and Jan Born. Sleep in children improves memory performance on declarative but not procedural tasks. Learning & memory, 15(5):373–377, 2008.

[63] W Hager and M Hasselhorn. Handbuch deutschsprachiger wortnormen hogrefe. Verlag für Psychologie Göttingen, Bern, Toronto, Seattle, 1994.

[64] Björn Rasch, Christian Büchel, Steffen Gais, and Jan Born. Odor cues during slow-wave sleep prompt declarative memory consolidation. Science, 315(5817):1426– 1429, 2007.

[65] Peter Zimmermann and Bruno Fimm. Testbatterie zur Aufmerksamkeitsprüfung:(TAP), Version 1.7. Psytest, 2002.

[66] Maria Laura Ferster, Caroline Lustenberger, and Walter Karlen. Configurable mobile system for autonomous high-quality sleep monitoring and closed-loop acoustic stimulation. IEEE Sensors Letters, 3(5):1–4, 2019.

[67] Arnaud Delorme and Scott Makeig. Eeglab: an open source toolbox for analysis of single-trial eeg dynamics including independent component analysis. Journal of neuroscience methods, 134(1):9–21, 2004.

[68] Richard B Berry, Rita Brooks, Charlene Gamaldo, Susan M Harding, Robin M Lloyd, Stuart F Quan, Matthew T Troester, and Bradley V Vaughn. Aasm scoring manual updates for 2017 (version 2.4), 2017.

[69] Sven Leach, Georgia Sousouri, and Reto Huber. ‘high-density-sleepcleaner’: An open-source, semi-automatic artifact removal routine tailored to high-density sleep eeg. Journal of Neuroscience Methods, 391:109849, 2023.

[70] Reto Huber, M Felice Ghilardi, Marcello Massimini, and Giulio Tononi. Local sleep and learning. Nature, 430(6995):78–81, 2004.

[71] Philipp Berens. Circstat: a matlab toolbox for circular statistics. Journal of statistical software, 31:1–21, 2009.

[72] Mike X Cohen. Analyzing neural time series data: theory and practice. MIT press, 2014.

[73] Scott M Doran, Hans PA Van Dongen, and David F Dinges. Sustained attention performance during sleep deprivation: evidence of state instability. Archives italiennes de biologie, 139(3):253–267, 2001.

[74] Glenn Gunzelmann, Joshua B Gross, Kevin A Gluck, and David F Dinges. Sleep deprivation and sustained attention performance: Integrating mathematical and cognitive modeling. Cognitive science, 33(5):880–910, 2009.

[75] Thomas E Nichols and Andrew P Holmes. Nonparametric permutation tests for functional neuroimaging: a primer with examples. Human brain mapping, 15(1): 1–25, 2002.

[76] Yoav Benjamini and Yosef Hochberg. Controlling the false discovery rate: a practical and powerful approach to multiple testing. Journal of the Royal statistical society: series B (Methodological), 57(1):289–300, 1995.

