## Supplement for "Acoustically evoked K-complexes together with sleep spindles boost verbal declarative memory consolidation in healthy adults"

### Supplemental information

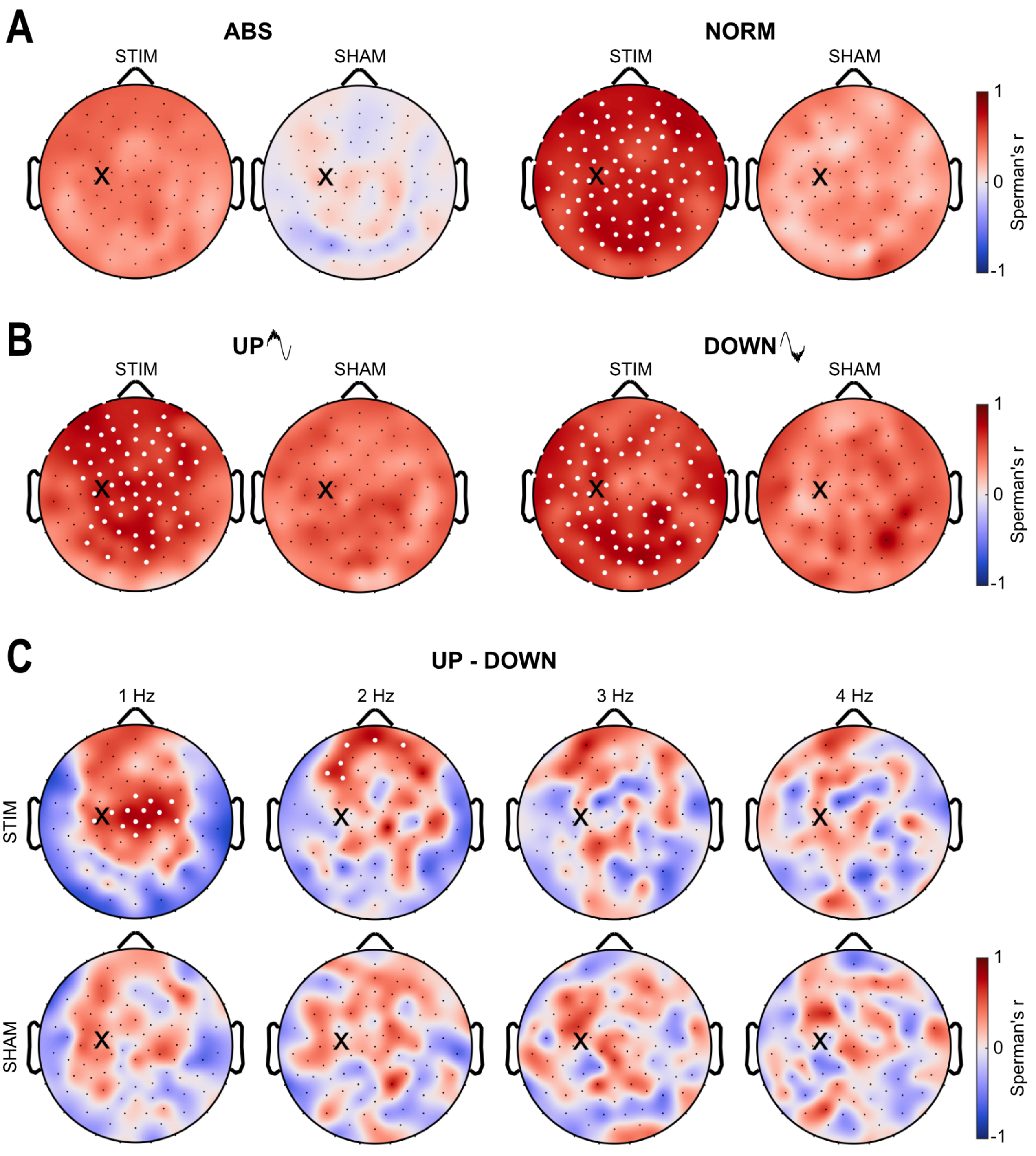


**Supplementary Fig. 1. Topography of Spearman's rank correlation coefficient between overnight verbal memory performance and stimulus-associated sleep spindles, seperate for STIM and SHAM. A)** Correlation with the absolute number of spindles (left) and the normalized number of spindles (right). Normalization was performed by expressing the absolute number of stimulus-associated spindles relative to the total number of overnight spindles. **B)** Correlation with the normalized number of stimulus-evoked sleep spindles coupled to the up-phase (left) and the down-phase (right) of the underlying 1 Hz wave. **C)** Correlation of the difference in the normalized number of spindles coupled to the up- and down-phase of the underlying slow wave, separated by slow wave frequencies. White dots represent electrodes indicating significant correlations with overnight verbal memory performance (Spearman's rank test; $p < .05$; cluster corrected); the black cross indicates the target electrode.

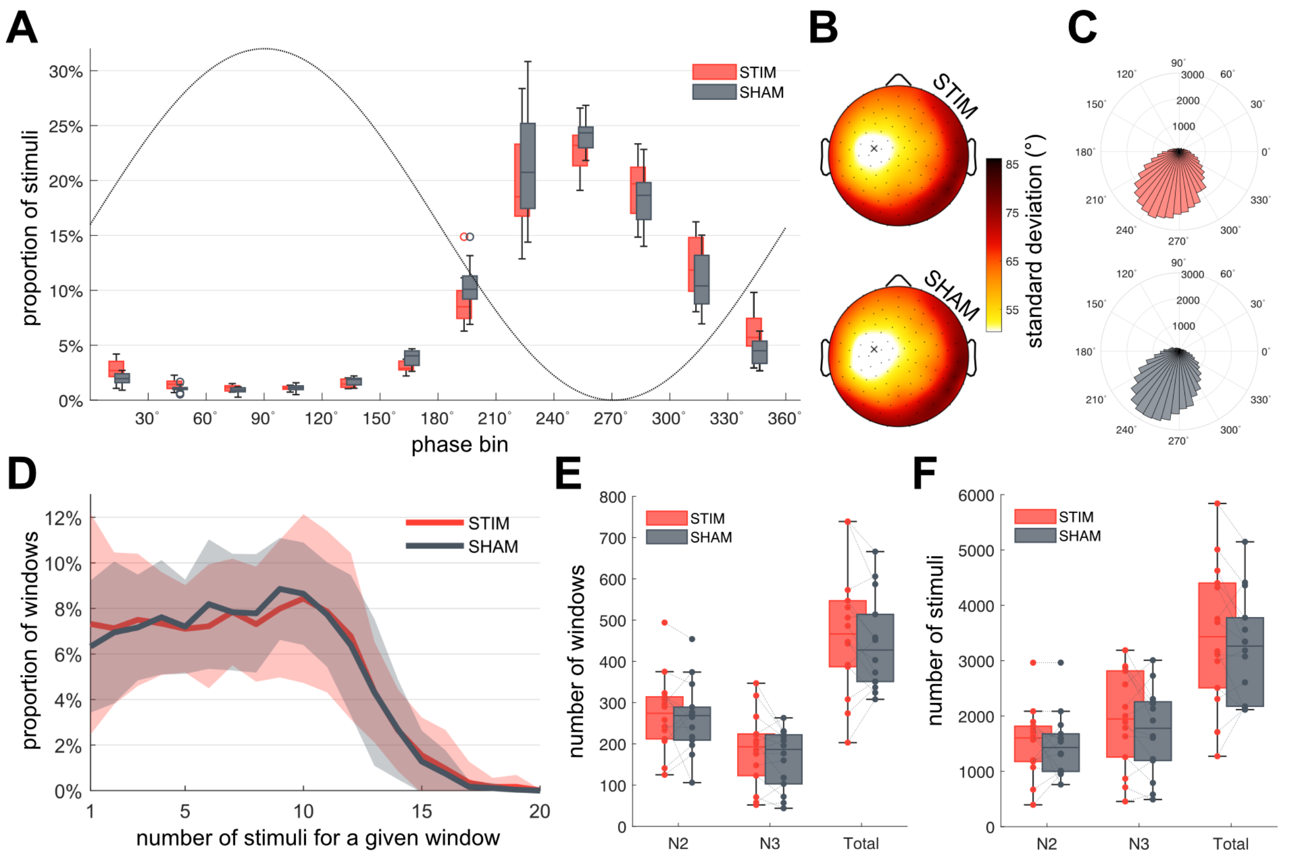


**Supplementary Fig. 2. Stimulation details.** All figures display stimulation details for the night with (red) and without stimulation (gray). **A)** Proportion of stimuli falling into a specific phase bin (30°steps). Proportions were calculated individually for each participant. Boxplots illustrate the distribution among participants. **B)** Topographic phase precision expressed as the circular standard deviation of phase values at times of stimulation. For a given electrode, the circular standard deviation was calculated individually per participant. Then, for a given electrode, those values were averaged (circular mean) over participants. **C)** Phase distribution of all stimuli, pooled over participants. **D)** Proportion of windows containing a certain number of stimuli, calculated individually per participant. The mean±SD over participants is displayed. **E)** Total number of ON|OFF window-pairs falling into an N2 or N3 epoch, as well as the total number of windows. Window-pairs were assigned to the sleep stage the majority of the window-pair belonged to. Each dot represents the value from a single participant. **F)** Total number of stimuli in N2, N3, and in total. Each dot represents a single participant.

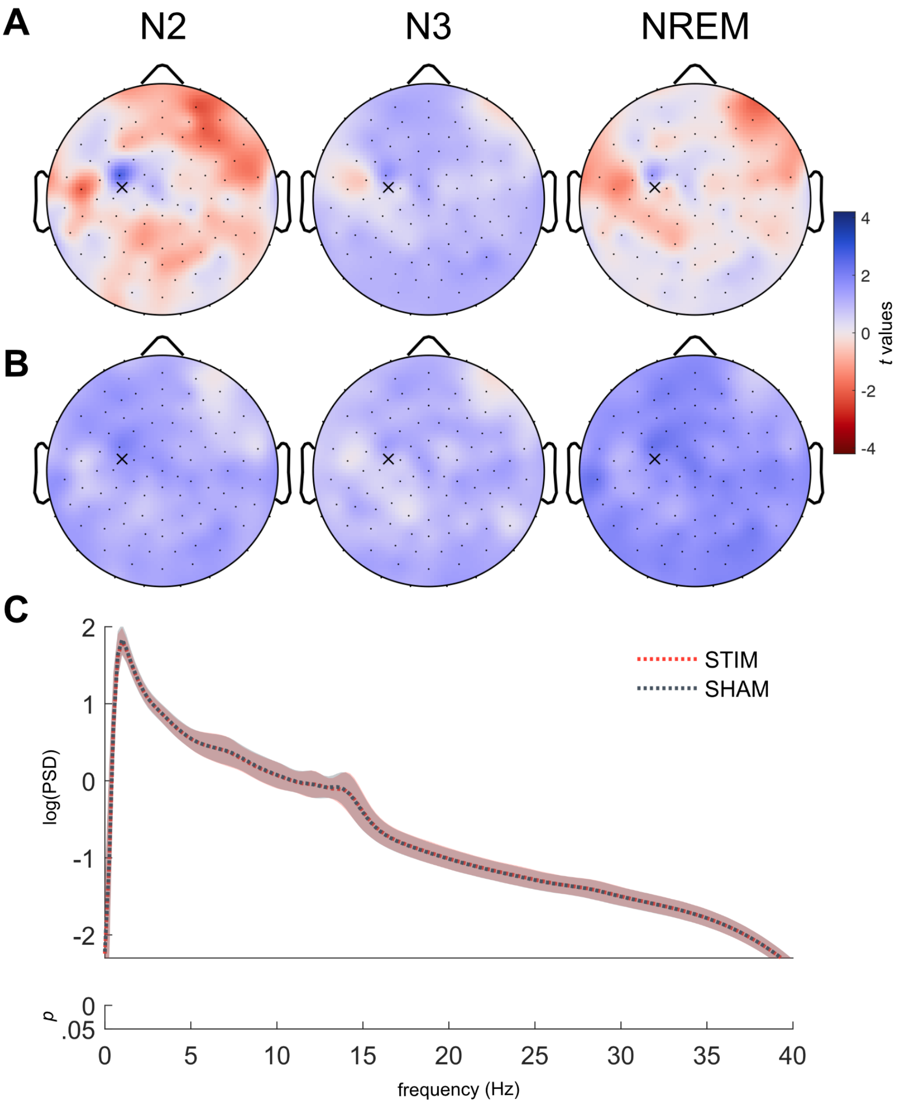


**D**

| | STIM | SHAM | *t*(13) | p value |
|---|---|---|---|---|
| **TST (min)** | 430.48 (44.49) | 432.31 (39.30) | -0.187 | .855 |
| **SL (min)** | 18.48 (8.88) | 22.98 (15.94) | -1.469 | .166 |
| **NREM (min)** | 337.33 (26.52) | 334.69 (27.36) | 0.328 | .748 |
| **N1 (min)** | 29.43 (11.79) | 29.50 (16.41) | -0.024 | .981 |
| **N2 (min)** | 216.10 (26.41) | 211.48 (29.40) | 0.686 | .505 |
| **N3 (min)** | 91.81 (34.12) | 93.71 (30.53) | -0.396 | .699 |
| **N2 + N3 (min)** | 307.90 (31.02) | 305.19 (31.92) | 0.328 | .748 |
| **REM (min)** | 93.14 (25.90) | 97.62 (27.51) | -0.836 | .418 |
| **WAKE (min)** | 47.76 (43.35) | 45.76 (39.26) | 0.212 | .835 |
| **WASO (min)** | 34.02 (43.55) | 27.83 (36.61) | 0.878 | .396 |
| **nWASO (#)** | 19.50 (6.04) | 16.50 (5.10) | 1.738 | .106 |
| **SEFF (%)** | 90.00 (9.09) | 90.43 (8.18) | -0.219 | .830 |

**Supplementary Fig. 3. Overnight analyses. A)** Topographic overnight slow-wave activity (SWA, 1 – 4.5 Hz) expressed as the difference in SWA between conditions (STIM− SHAM). Mean SWA was computed in N2 (left), N3 (middle) and NREM (N2+N3; right) epochs. Paired t-tests (cluster corrected) indicated no significant differences between conditions (all $p \leq .05$). **B)** Topographic relative overnight SWA. For a given participant, SWA levels were normalized by the channel average. Paired t-tests (cluster corrected) indicated no significant differences between conditions (all $p \leq .05$). **C)** Logarithmic power spectral density (PSD) from 0 to 40 Hz expressed as the average over all NREM epochs and channels. The mean$\pm$SD over participants is displayed. Paired t-tests (FDR corrected) indicated no significant differences between conditions (all $p \leq .05$). **D)** Sleep architecture. Paired *t*-test test statistics are reported (uncorrected). There are no indications for auditory stimulation affecting sleep parameters. Total sleep time (TST); sleep latency (SL); non-rapid eye movement sleep (NREM, stages N1 to N3); rapid eye movement sleep (REM); wake after sleep onset (WASO); number of awakenings after sleep onset (nWASO); sleep efficiency (SEFF).

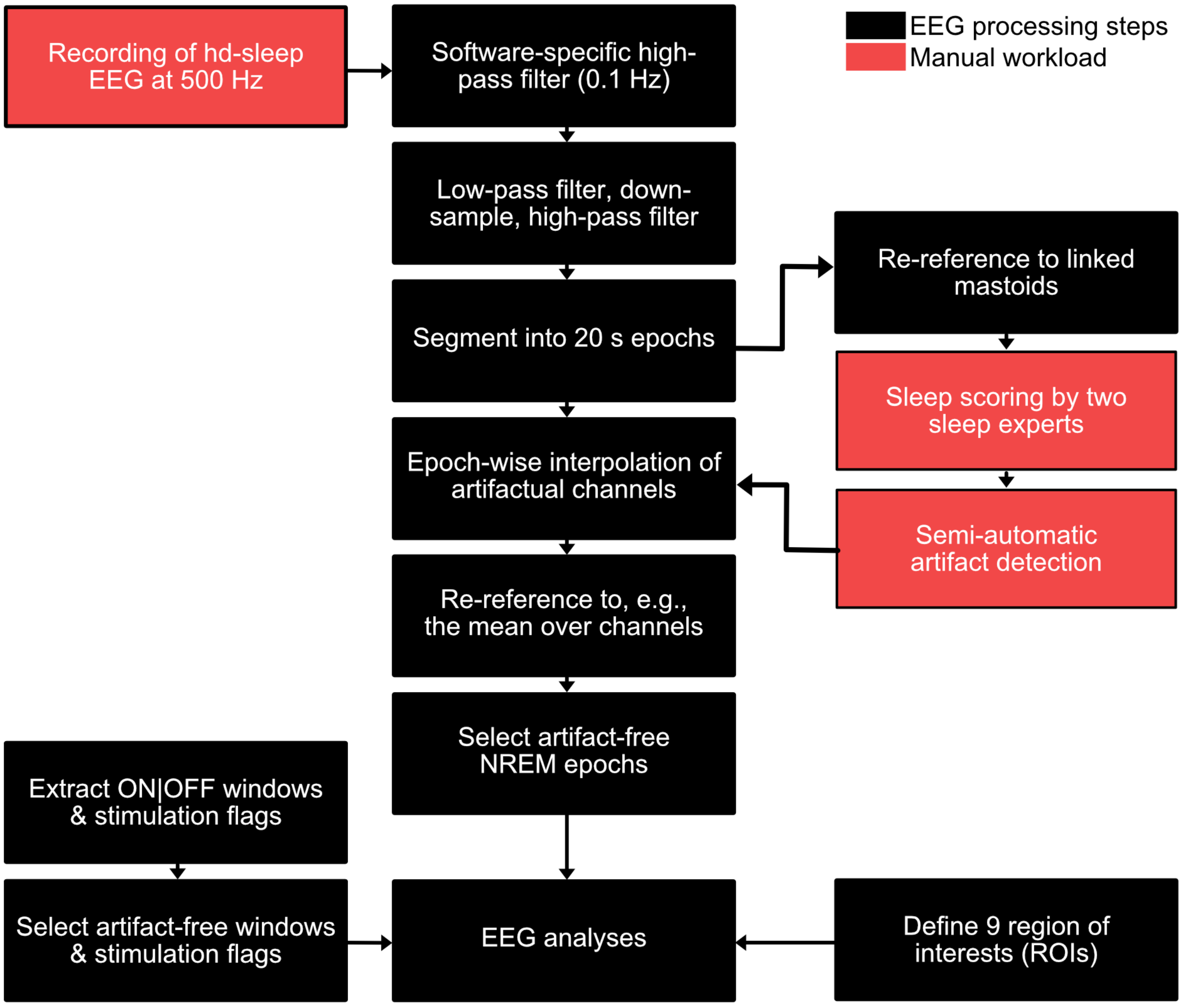


**Supplementary Fig. 4. Preprocessing pipeline.** The recorded EEG was exported with a soft-ware specific high-pass filter and imported into Matlab. The EEG was low-pass filtered, down-sampled, high-pass filtered, and segmented into 20 s epochs. Each night was sleep-scored by two sleep experts and artifacts were detected semi-automatically. Artifactual data of affected channels in affected epochs was interpolated using epoch-wise interpolation. Thereafter, the EEG was average referenced and NREM (N2+N3) epochs were selected for further analyses. If applicable, analyses were performed based on stimulations, stimulation windows, and/or in nine regions of interest (ROIs).

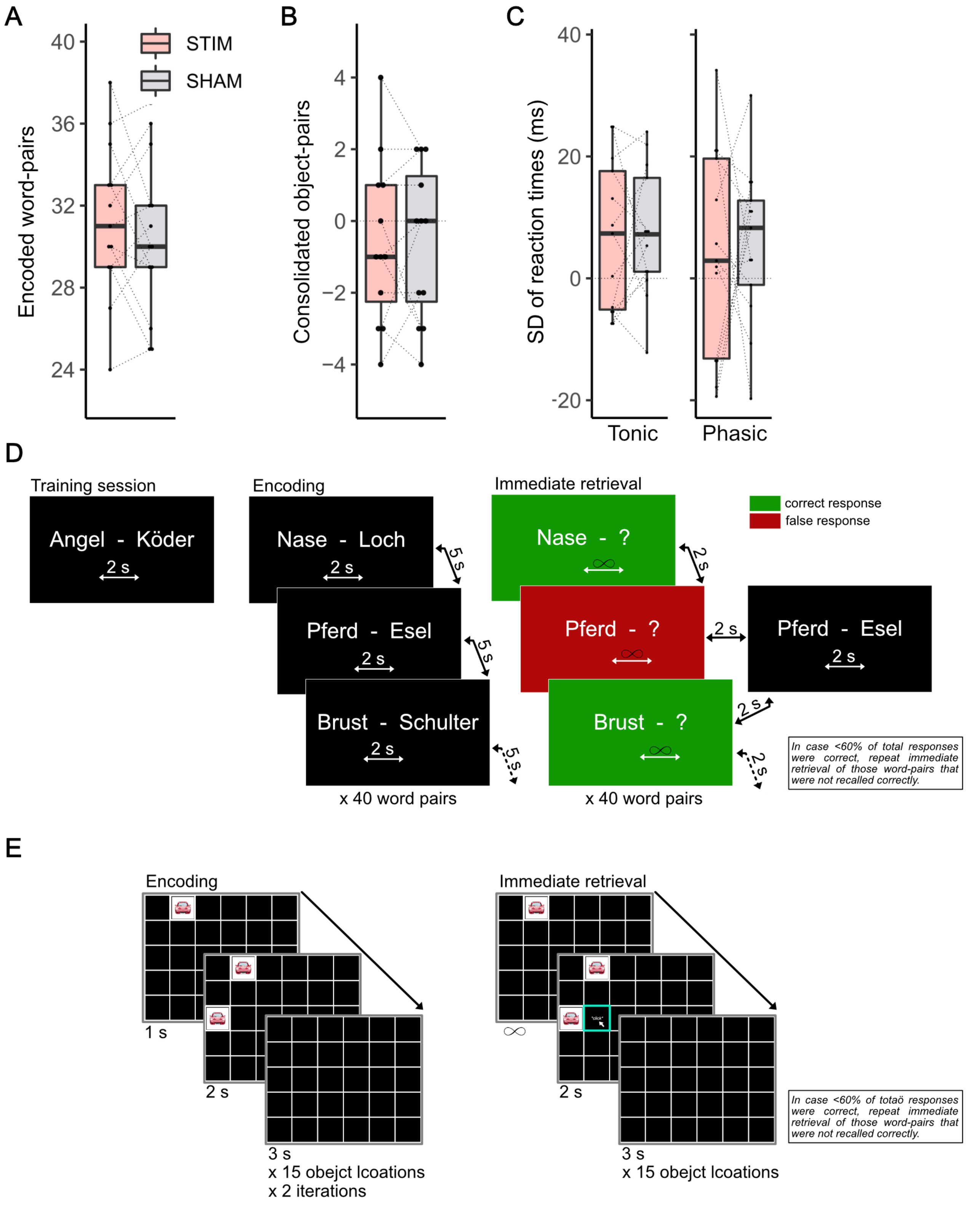


**Supplementary Fig. 5. Behavioral tests. A)** Retrieval performance of newly encoded word-pairs following a night with (STIM) and without auditory stimulation (SHAM). **B)** Overnight memory change of successfully remembered object-pairs (morning − evening). Dots represent the performance of single participants. **C)** Standard deviation of reaction times (morning − evening) assessing tonic (left) and phasic alertness (right). Tonic alertness is measured by presenting a cross on the monitor at randomly varying intervals to which the participant should respond as quickly as possible by pressing a key. Phasic alterness (temporal orientation of attentional focus) is measured by introducing a preceding warning tone before the cross is presented. **D)** Procedure of the associated word-pair memory task. After a training word-pair (left), an encoding phase followed during which participants learned 40 associated word-pairs (middle). Thereafter, participants immediately retrieved the newly encoded word-pairs (right). After a correct response, the next word-pair followed. After a wrong response, the whole word-pair was presented once more. The procedure was repeated until $\geq$60% of word-pairs were recalled correctly. Note that the screen was not visible for the participant, as word-pairs were presented orally. **E)** Procedure of the object-location task. During encoding (left), the location of 15 object-pairs was presented twice. Thereafter, participants immediately retrieved the newly encoded object-pairs (middle). The tile selected by participants was highlighted upon mouse-click in cyan.

#### List 1

| | | | | |
|---|---|---|---|---|
| 1 | Nase | Loch | Nose | Hole |
| 2 | Pferd | Esel | Horse | Donkey |
| 3 | Pulli | Kragen | Sweater | Collar |
| 4 | Raupe | Kohl | Caterpillar | Cabbage |
| 5 | Schaukelstuhl | Oma | Walker | Grandmother |
| 6 | Bikini | Strand | Bikini | Beach |
| 7 | Delphin | Robbe | Dolphin | Seal |
| 8 | Eule | Adler | Owl | Eagle |
| 9 | Frosch | Fliege | Frog | Fly |
| 10 | Gummistiefel | Jacke | Boots | Jacket |
| 11 | Mauer | Stein | Wall | Stone |
| 12 | Mantel | Handschuh | Coat | Glove |
| 13 | Krebs | Augen | Crab | Eyes |
| 14 | Seehund | Schwanz | Walrus | Tail |
| 15 | Sessel | Decke | Armchair | Blanket |
| 16 | Stier | Arena | Bull | Arena |
| 17 | Kueche | Salat | Kitchen | Salad |
| 18 | Tuer | Keller | Door | Cellar |
| 19 | Uhr | Diamant | Watch | Diamond |
| 20 | Erde | Kartoffel | Earth | Potato |
| 21 | Gras | Schlange | Grass | Snake |
| 22 | Wald | Fuchs | Forest | Fox |
| 23 | Bluete | Schmetterling | Blossom | Butterfly |
| 24 | Auftrag | Arbeit | Assignment | Work |
| 25 | Brust | Schulter | Chest | Shoulder |
| 26 | Stirn | Muetze | Forehead | Hat |
| 27 | Strumpf | Stoff | Sock | Fabric |
| 28 | Fass | Schiff | Barrel | Ship |
| 29 | Geschoss | Kugel | Projectile | Bullet |
| 30 | Sumpf | Schlamm | Swamp | Mud |
| 31 | Huette | Tal | Cottage | Valley |
| 32 | Halle | Palast | Hall | Palace |
| 33 | Ereignis | Fest | Event | Festival |
| 34 | Glaube | Verzicht | Belief | Abstinence |
| 35 | Herrscher | Befehl | Ruler | Order |
| 36 | Flocken | Bergung | Flake | Rescue |
| 37 | Brechstange | Schloss | Crowbar | Lock |
| 38 | Allee | Dickicht | Road | Bushes |
| 39 | Angabe | Zeuge | Statement | Witness |
| 40 | Aufstand | Schild | Riot | Shield |

#### List 3

| | | | | |
|---|---|---|---|---|
| 1 | Truhe | Gold | Treasure | Gold |
| 2 | Vogel | Rabe | Bird | Raven |
| 3 | Waschbecken | Zahnbuerste | Sink | Toothbrush |
| 4 | Zeigefinger | Nagel | Finger | Nail |
| 5 | Garage | Haus | Garage | House |
| 6 | Tasse | Loeffel | Mug | Spoon |
| 7 | Telefon | Freund | Telephone | Friend |
| 8 | Erdbeere | Kuchen | Strawberry | Cake |
| 9 | Schule | Tafel | School | Blackboard |
| 10 | Urlaub | Sonne | Vacation | Sun |
| 11 | Geld | Muenze | Money | Coin |
| 12 | Lied | Text | Song | Text |
| 13 | Koffer | Reise | Suitcase | Journey |
| 14 | Zahn | Mund | Tooth | Mouth |
| 15 | Schnee | Berg | Snow | Mountain |
| 16 | Maus | Kaese | Mouse | Cheese |
| 17 | Papier | Brief | Sheet | Letter |
| 18 | Zeitung | Bleistift | Newspaper | Pencil |
| 19 | Weste | Knopf | Vest | Button |
| 20 | Zehen | Knie | Toe | Knee |
| 21 | Augenbraue | Pupille | Eyebrow | Pupil |
| 22 | Pflanze | Getreide | Plant | Grain |
| 23 | Fischer | See | Fisherman | Lake |
| 24 | Einbrecher | Polizist | Burglar | Police |
| 25 | Kind | Puppe | Child | Doll |
| 26 | Lager | Raeuber | Camp | Bandit |
| 27 | Gletscher | Felsblock | Glacier | Boulder |
| 28 | Zucker | Apfel | Sugar | Apple |
| 29 | Koerper | Gelenk | Body | Joint |
| 30 | Schatten | Scheinwerfer | Shadow | Spotlight |
| 31 | Feuer | Stern | Fire | Star |
| 32 | Klippe | Lawine | Cliff | Avalanche |
| 33 | Riese | Schritt | Giant | Step |
| 34 | Sport | Zeit | Sport | Time |
| 35 | Museum | Fund | Museum | Discovery |
| 36 | Ufer | Damm | Shore | Dam |
| 37 | Anstand | Hoeflichkeit | Manners | Politeness |
| 38 | Aquarell | Galerie | Watercolor | Gallery |
| 39 | Beruf | Fleischer | Job | Butcher |
| 40 | Bibliothek | Signatur | Library | Signature |

#### List 2

| | | | | |
|---|---|---|---|---|
| 1 | Loewe | Zoo | Lion | Zoo |
| 2 | Ofen | Brot | Oven | Bread |
| 3 | Ohr | Backe | Ear | Cheek |
| 4 | Ampel | Auto | Signal | Car |
| 5 | Saege | Axt | Saw | Axe |
| 6 | Schal | Kaelte | Scarf | Cold |
| 7 | Schaufel | Garten | Shovel | Garden |
| 8 | Schere | Blatt | Scissors | Sheet |
| 9 | Arm | Blut | Arm | Blood |
| 10 | Badewanne | Ente | Bathtub | Duck |
| 11 | Bart | Haut | Beard | Skin |
| 12 | Bauch | Herz | Belly | Heart |
| 13 | Bein | Knochen | Leg | Bone |
| 14 | Bett | Schlaf | Bed | Sleep |
| 15 | Daumen | Ringfinger | Thumb | Hand |
| 16 | Hund | Dackel | Dog | Leash |
| 17 | Post | Fahrrad | Post | Bicycle |
| 18 | Kirche | Glocken | Church | Bells |
| 19 | Heft | Note | Notebook | Grade |
| 20 | Zimmer | Hotel | Room | Hotel |
| 21 | Butter | Kuehlschrank | Butter | Fridge |
| 22 | Becher | Pudding | Cup | Pudding |
| 23 | Kaffee | Dampf | Coffee | Steam |
| 24 | Gewitter | Wasser | Storm | Water |
| 25 | Guertel | Leder | Belt | Leather |
| 26 | Hammer | Zange | Hammer | Pliers |
| 27 | Schrank | Griff | Closet | Handle |
| 28 | Stuhl | Polster | Chair | Cushion |
| 29 | Harfe | Klavier | Harp | Piano |
| 30 | Kueste | Meer | Coast | Sea |
| 31 | Tinte | Plakat | Ink | Poster |
| 32 | Gitter | Gefaengnis | Bars | Prison |
| 33 | Reptil | Insekt | Reptile | Insect |
| 34 | Zitrone | Pfirsich | Lemon | Peach |
| 35 | Zigarre | Pfeife | Cigar | Pipe |
| 36 | Infektion | Schmerzen | Infection | Pain |
| 37 | Instrument | Oboe | Instrument | Oboe |
| 38 | Paechter | Vertrag | Tenant | Contract |
| 39 | Schauspiel | Ausdruck | Play | Expression |
| 40 | Stift | Kappe | Pen | Cover |

#### List 4

| | | | | |
|---|---|---|---|---|
| 1 | Buch | Brille | Book | Glasses |
| 2 | Strasse | Laterne | Street | Lantern |
| 3 | Heizung | Winter | Heating | Winter |
| 4 | Gitarre | Konzert | Guitar | Concert |
| 5 | Foto | Rahmen | Photo | Frame |
| 6 | Radio | Stimme | Radio | Voice |
| 7 | Baum | Natur | Tree | Nature |
| 8 | Fenster | Wetter | Window | Weather |
| 9 | Regal | Brett | Shelf | Board |
| 10 | Gemuese | Kochtopf | Vegetable | Pot |
| 11 | Kerze | Nacht | Candle | Night |
| 12 | Teppich | Flug | Carpet | Flight |
| 13 | Parkplatz | Einkauf | Parking | Shopping |
| 14 | Katze | Fell | Cat | Fur |
| 15 | Kino | Leinwand | Cinema | Screen |
| 16 | Himmel | Gebet | Heaven | Prayer |
| 17 | Getraenk | Schaum | Drink | Foam |
| 18 | Schueler | Buecherei | Student | Bookstore |
| 19 | Doktor | Krankenhaus | Doctor | Hospital |
| 20 | Maschine | Eisenbahn | Machine | Train |
| 21 | Messer | Metall | Knife | Metal |
| 22 | Flasche | Korken | Bottle | Cork |
| 23 | Lampe | Schirm | Lamp | Shade |
| 24 | Giesskanne | Rost | Chain | Rust |
| 25 | Panne | Anruf | Emergency | Call |
| 26 | Streichholz | Fabrik | Match | Factory |
| 27 | Bakterien | Mikroskop | Bacteria | Microscope |
| 28 | Kleidung | Buegeleisen | Clothing | Iron |
| 29 | Staub | Dachboden | Dust | Attic |
| 30 | Verkaeufer | Moebel | Vendor | Furniture |
| 31 | Eisen | Schmied | Furnace | Smith |
| 32 | Naesse | Fluss | Current | River |
| 33 | Laden | Reklame | Store | Promotion |
| 34 | Landschaft | Moor | Landscape | Moor |
| 35 | Macht | Kampf | Power | Fight |
| 36 | Labor | Pipette | Laboratory | Pipette |
| 37 | Schlinge | Seil | Loop | Rope |
| 38 | Maedchen | Verlobung | Girl | Engagement |
| 39 | Naht | Kreuzstich | Seam | Needle |
| 40 | Orkan | Luft | Hurricane | Air |

**Supplementary Fig. 6. Wordlists used in the associated word-pair memory task.**

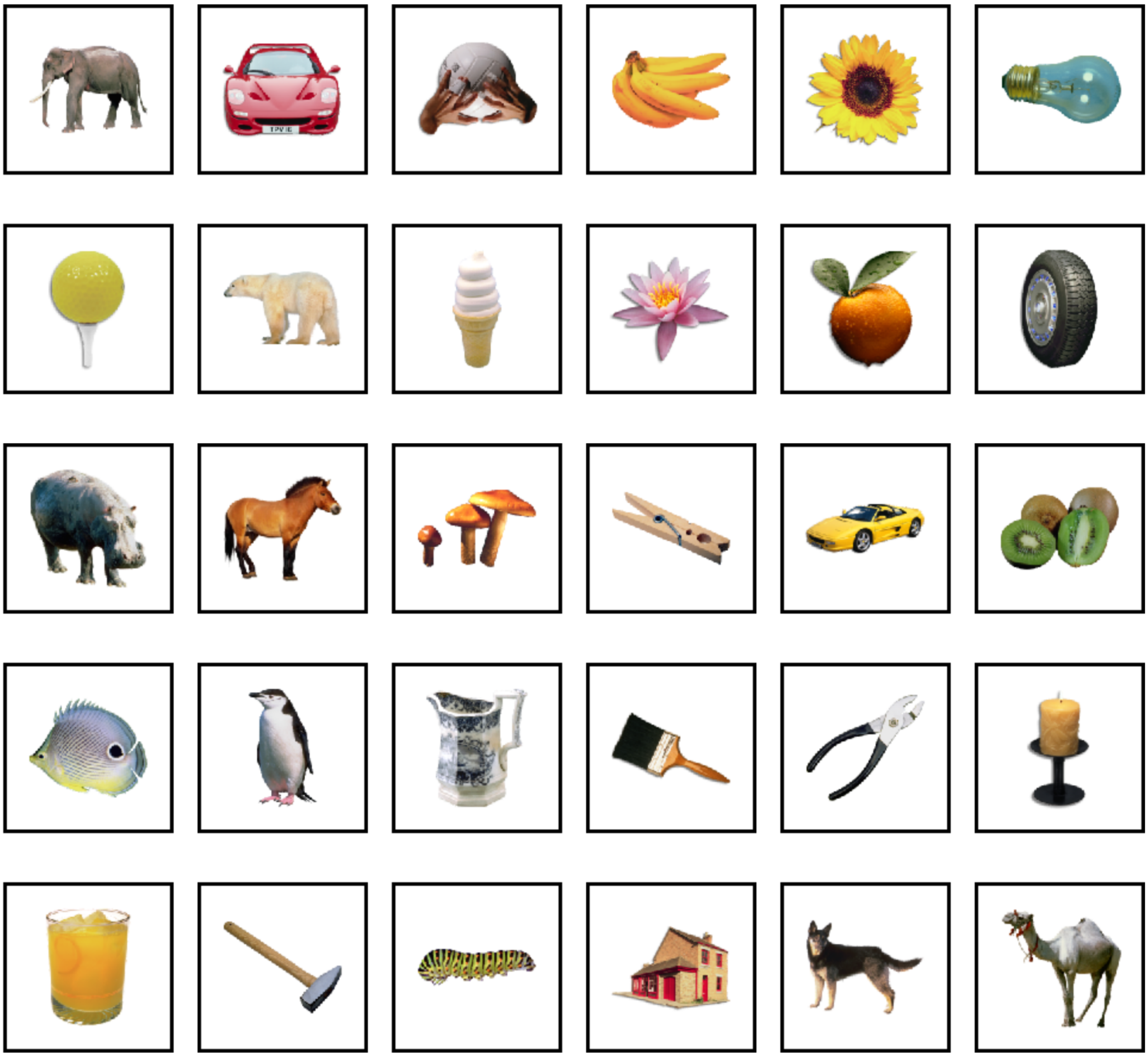

**Supplementary Fig. 7. Objects of object-location task.**

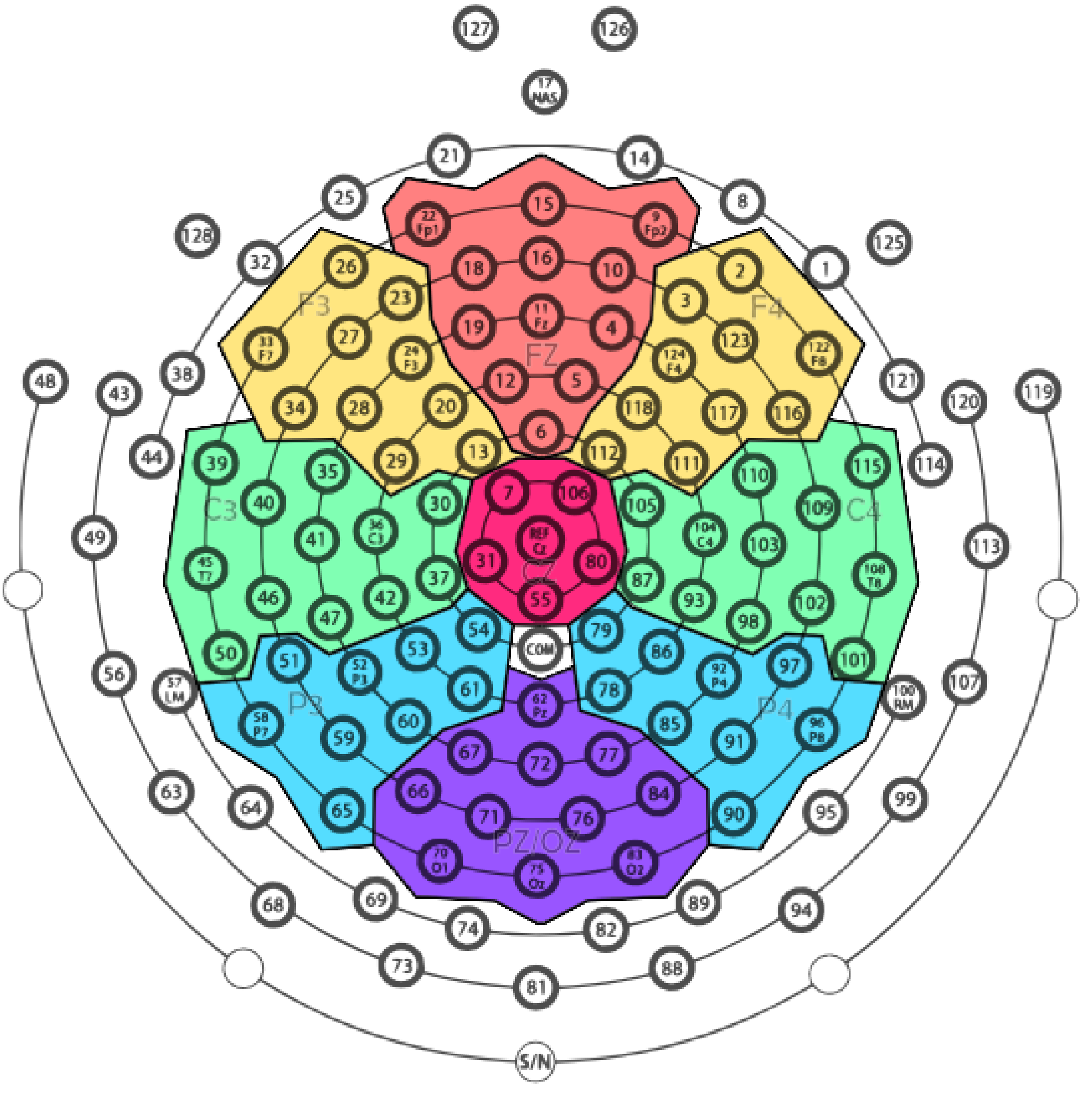


**Supplementary Fig. 8. Regions of interest.** Colors indicate channels that were included in regions of interest (ROIs) analyses. Respective 10-20 electrodes within ROIs (Fz, F3, F4, etc.) were used to refer to the respective ROI.
